# The murine meninges acquire lymphoid tissue properties and harbour autoreactive B cells during chronic *Trypanosoma brucei* infection

**DOI:** 10.1101/2023.04.30.538710

**Authors:** Juan F. Quintana, Matthew C. Sinton, Praveena Chandrasegaran, Lalit Kumar Dubey, John Ogunsola, Moumen Al Samman, Michael Haley, Gail McConnell, Nono-Raymond Kuispond Swar, Dieudonne Mumba Ngoyi, David Bending, Luis de Lecea, Annette MacLeod, Neil A. Mabbott

## Abstract

The meningeal space is a critical brain structure providing immunosurveillance for the central nervous system, but the impact of infections on the meningeal immune landscape is far from being fully understood. The extracellular protozoan parasite *Trypanosoma brucei*, which causes Human African Trypanosomiasis (HAT) or sleeping sickness, accumulates in the meningeal spaces, ultimately inducing severe meningitis and resulting in death if left untreated. Thus, sleeping sickness represents an attractive model to study immunological dynamics in the meninges during infection. Here, by combining single cell transcriptomics and mass cytometry by time of flight (CyTOF) with *in vivo* interventions, we found that chronic *T. brucei* infection triggers the development of ectopic lymphoid aggregates (ELAs) in the murine meninges. These infection-induced ELAs were defined by the presence of ER-TR7^+^ fibroblastic reticular cells, CD21/35^+^ follicular dendritic cells, CXCR5^+^ PD1^+^ T follicular helper-like phenotype, GL7^+^ CD95^+^ GC-like B cells, and plasmablasts/plasma cells. Furthermore, the B cells found in the infected meninges produced high-affinity autoantibodies able to recognise mouse brain antigens, in a process dependent on LTβ signalling. A mid-throughput screening identified several host factors recognised by these autoantibodies, including myelin basic protein (MBP), coinciding with cortical demyelination and brain pathology. In humans, we identified the presence of autoreactive IgG antibodies in the cerebrospinal fluid of second stage HAT patients that recognised human brain lysates and MBP, consistent with our findings in experimental infections. Lastly, we found that the pathological B cell responses we observed in the meninges required the presence of *T. brucei* in the CNS, as suramin treatment before the onset of the CNS stage prevented the accumulation of GL7^+^ CD95^+^ GC-like B cells and brain-specific autoantibody deposition. Taken together, our data provide evidence that the meningeal immune response during chronic *T. brucei* infection results in the acquisition of lymphoid tissue-like properties, broadening our understanding of meningeal immunity in the context of chronic infections. These findings have wider implications for understanding the mechanisms underlying the formation ELAs during chronic inflammation resulting in autoimmunity in mice and humans, as observed in other autoimmune neurodegenerative disorders, including neuropsychiatric lupus and multiple sclerosis.

## Introduction

The meningeal space is rapidly being recognised as a critical site for immunological responses in the central nervous system (CNS) under homeostasis^1–3^, aging^4^, and as a consequence of insults such as traumatic brain injury^5^ and infection^6–8^. The extracellular protozoan parasite *Trypanosoma brucei*, which causes human African trypanosomiasis (HAT; sleeping sickness in humans) and animal African trypanosomiasis (Nagana in domestic animals) accumulates in the CNS and meningeal spaces triggering severe meningitis^9,10^. This culminates in the development of a wide range of debilitating neurological disorders^9,11–13^. These symptoms are diverse and include fatigue, altered sleep and circadian patterns, tremors, motor weakness, epilepsy, paralysis of one or more extremities, and Parkinson-like abnormal body movements^14–16^. Consistent with clinical data from humans, experimental trypanosomiasis in mice also results in chronic infection, leading to altered behaviour^17–20^. Thus, murine infection with *T. brucei* is a useful model to investigate meningeal responses to infection.

Chronic inflammatory processes are known to result in the formation of ectopic lymphoid aggregates (ELAs)^21–24^. Indeed, ELAs have been reported in a wide range of autoimmune disorders, including those affecting the CNS such as neuropsychiatric lupus^25^ and multiple sclerosis^26^. The diverse cytokine and chemokine repertoire found in chronically inflamed tissues, including lymphotoxin-α/-β (LTα and LTβ) and CXCL13, help to the create interactive niches needed to generate such structures^21–24^. Stromal LTβ receptor (LTβR) signalling is important in generating the microarchitecture required for efficient antigen presentation and follicle organisation, which typically includes collagen-rich reticular cords that serve as channels for cellular trafficking, immunological synapses, and B cell affinity maturation^27^. Similarly, CXCL13 is an important chemokine for defining local gradients controlling B cell domains, typically in proximity to follicular dendritic cells (FDCs) and CD4^+^ T follicular helper cells (T_FH_) inducing the formation of germinal centres (GC), in which B cells undergo affinity maturation and somatic hypermutation to generate high affinity antibodies^21,23,25^. These reactions are typically restricted to secondary lymphoid organs such as the spleen and lymph nodes, but can occur ectopically in response to chronic inflammation, and may result in pathological consequences such as the formation of autoreactive antibodies, as recently described for multiple sclerosis and neuropsychiatric lupus^23,25,26,28^.

In secondary lymphoid organs, including the spleen and lymph nodes, lymphatic vessels act as conduits for the transport of tissue-derived antigens and dendritic cells to lymph nodes, where naïve and memory T cells are optimally positioned for the detection of their cognate antigen^29–31^. Similarly, immune complexes can be acquired by macrophages in the subcapular space in lymph nodes and transferred directly to FDCs and B cells^32^. However, several key findings in recent years have led to a better understanding of the role of lymphatic vessels in the dura mater layer of the meninges. For instance, the meningeal lymphatic vessels can convey macromolecular complexes and immune cells from the meninges and cerebrospinal fluid (CSF) to the deep cervical lymph nodes^4,33,34^. However, it is also plausible that extramedullary reactions may take place locally in the meningeal spaces and brain borders, as reported recently in neuropsychiatric lupus^25^ and multiple sclerosism^28^. Whether the same extramedullar immunological reactions in the brain and/or meninges, reminiscent of those taking place in secondary lymphoid tissues, can be triggered by chronic, unresolved infections is uncertain.

Here, we investigated how the meningeal transcriptional environment is altered during *T. brucei* infection, using a combination of single cell transcriptomics and mass cytometry by time-of-flight (CyTOF). We found that chronic *T. brucei* infection in the meningeal space results in a broad rearrangement of the immune landscape in the murine meninges, with a significant increase in the frequency of innate (mononuclear phagocytes and granulocytes) and adaptive (T, NKT, and B cells) immune cells. Furthermore, we identified a population of autoreactive B cells in the meningeal spaces, including the leptomeninges. These autoreactive B cels were able to recognise mouse brain antigens and deposit high-affinity IgG antibodies in several brain areas including the hippocampus and cortex, and this deposition was associated with cortical and white mater demyelination. We also detected significant levels of autoreactive IgM and IgG antibodies in the cerebrospinal fluid of HAT patients with inflammatory encephalopathy. Furthermore, using a targeted screening approach we identified myelin basic protein as one of the host antigens detected by autoreactive IgG antibodies in mouse serum and human CSF collected during the chronic stage of the infection. Taken together, this study demonstrates that the meningeal landscape acquires lymphoid tissue-like properties resulting in the formation of autoreactive B cells. These results imply that the chronic brain inflammation induced by African trypanosomes results in an autoimmune disorder affecting the brain as observed in other neurological disorders of unknown aetiology such as neuropsychiatric lupus and multiple sclerosis. We anticipate that the data presented here will pave the way to understanding how chronic meningitis results in impaired peripheral tolerance and the development of autoimmunity in the context of chronic infections.

## Materials and methods

### Ethical statement

All animal experiments were approved by the University of Glasgow Ethical Review Committee and performed in accordance with the Home Office guidelines, UK Animals (Scientific Procedures) Act, 1986 and EU directive 2010/63/EU. All experiments were conducted under SAPO regulations and UK Home Office project licence number PP4863348 to Annette Macleod. The *in vivo* work presented in this study was conducted at 30-days post-infection (dpi) and correlated with increased clinical scores and procedural severity. The archived human CSF samples from gambiense HAT patients from North Uganda used in this study were collected by Professor Wendy Bailey (Liverpool School of Tropical Medicine, UK). Ethical approval was given to Prof. Bailey by the Liverpool School of Tropical Medicine, UK, for sample collection and patients were provided with a written consent. We received ethical approved by the University of Glasgow MVLS Ethics Committee for Non-Clinical Research Involving Human Subjects (Reference no. 200120043) for the use of human archived samples. The CSF from healthy donors was obtained from the University of Edinburgh Brain and Tissue Bank and received ethical approval from the University of Edinburgh (REC 21/ES/0087).

### Murine infections with *Trypanosoma brucei*

Six-to eight-week-old female C57BL/6J mice (JAX, stock 000664) and C57BL/6-Tg(Nr4a1-EGFP/Cre)820Khog/J strain, also known as Nur77^GFP^ reporter mice (JAX, stock 016617), or the Nur77^Tempo^ reporter mouse line (kindly provided by Dr. David Bending), were inoculated by intra-peritoneal injection with ∼2 x 10^3^ parasites of strain *T. brucei brucei* Antat 1.1E^35^. Parasitaemia was monitored by regular sampling from tail venesection and examined using phase microscopy and the rapid “matching” method^36^. Uninfected mice of the same strain, sex and age served as uninfected controls. Mice were fed *ad libitum* and kept on a 12 h light– dark cycle. All the experiments were conducted between 8h and 12h. When using the *Nur77*^GFP^ or the *Nur77*^Tempo^ reporter mice, sample acquisition and analysis was conducted without *ex vivo* stimulation to preserve the TCR-dependent fluorescent reporter signal found in the tissue. For sample collection, we focussed on 30 days post-infection, as this has previously been shown to correlate with parasite infiltration in the epidural space^10,11^. Culture-adapted *T. brucei* Antat 1.1E whole cell lysates were prepared as followed. Parasites were cultured in HMI-9 supplemented with 10% FBS and 1% Penicillin/Streptomycin were grown at 37°C and 5% CO_2_ and harvested during the log phase. The parasites were harvested by centrifugation (800 g for 10 min at 4°C), washed three times in 1X PBS (Gibco) supplemented with cOmplete protease Inhibitor cocktail (Roche), and sonicated with 5 pulses of 10 seconds each. The resulting lysate was cleared by centrifugation (3,000g for 10 min at 4°C to remove cell debris), and the protein concentration of the cleared supernatant was measured using the Qubit protein kit (Thermo) and kept at –80°C until usage for ELISPOT and ELISA. For LTBR-Ig treatment, mice were inoculated with 1μg/μL of either LTBR-Ig or IgG2a i.p. (100 μl/mouse) for four consecutive days prior to infection, and then every seven days post-infection until culling. Preparation of single cell suspension from skull meninges for single-cell RNA sequencing

### Tissue processing and preparation of single cell suspension

Single-cell dissociations for scRNAseq experiments were performed as follow. Animals were infected for 30 days (*n* = 2 mice/pool, 2 independent pools per experimental condition), after which skullcap meninges were harvested for preparation of single cell suspensions. Uninfected animals were also included as naive controls (*n* = 3 mice/pool, 2 pools analysed). Briefly, all mice were killed by rapid decapitation following isoflurane anaesthesia, within the same time (between 7:00 and 9:00 AM). To discriminate circulating versus brain-resident immune cells, we performed intravascular staining of peripheral CD45^+^ immune cells, as previously reported^37^. Briefly, a total of 2 mg of anti-CD45-PE antibody (in 100 ml of 1X PBS) was injected intravenously 3 minutes prior culling. Mice were euthanised as described above and transcardially perfused with ice-cold 0.025% (wt/vol) EDTA in 1X PBS. The excised meninges were enzymatically digested with Collagenase P (1 mg/ml) and DNAse I (1 mg/ml; Sigma) in 1X PBS (HSBB) (Invitrogen) for ∼30 min at 37 °C. Single-cell suspensions were passed through 70 μm nylon mesh filters to remove any cell aggregates, and the circulating CD45-PE^+^ cells were removed from the single cell suspension using magnetic sorting with anti-PE microbeads (Miltenyi Biotec) according to manufacturer’s recommendations (**S1A Figure**).

## Mass cytometry sample processing

Single cell suspension from meninges were prepared as described above and resuspended in Dubecco’s Modified Eagle Medium (DMEM) to a concentration of 1 x 10^6^ cells/mL. Cells were activated for 6 h in a round-bottom 96-well plate using Cell Activation Cocktail (containing with Brefeldin A) (BioLegend, San Diego, USA) as per the manufacturer’s recommendations. Plates were then centrifuged at 300 x g for 5 min and the pellets resuspended in 50 µL of Cell-ID™ Cisplatin-195Pt viability reagent (Standard BioTools, San Francisco, USA), and incubated at room temperature for 2 min. Cells were washed twice in Maxpar® Cell Staining Buffer (Standard BioTools, San Francisco, USA), and centrifuged at 300 x g at room temperature for 5 min. The CD16/CD32 receptors were then blocked by incubating with a 1/50 dilution of TruStain FcX™ (BioLegend, San Diego, USA) in PBS at room temperature for 15 min. An antibody cocktail was prepared from the Maxpar® Mouse Sp/LN Phenotyping Panel Kit (Standard BioTools, San Francisco, USA), with and additional antibody against IgM. Cells were incubated with antibodies for 60 min, on ice before washing 3 times in Maxpar® Cell Staining Buffer (Standard BioTools, San Francisco, USA) as previously. Following staining, cells were fixed in 2% paraformaldehyde (PFA) overnight at 4°C. Cells were then washed twice with 1X eBioscience™ Permeabilization Buffer (Invitrogen, Waltham, USA) at 800 x g at room temperature for 5 min. The pellets were resuspended in intracellular antibody cocktail and incubated at room temperature for 45 min. Cells were washed 3 times in Maxpar® Cell Staining Buffer (Standard BioTools, San Francisco, USA) at 800 x g. The cells were then resuspended in 4% PFA at room temperature for 15 min, before collecting the cells at 800 x g and resuspending in Cell-ID™ Intercalator-Ir (Standard BioTools, San Francisco, USA). Finally, the cells were barcoded by transferring the stained cells to a fresh tube containing 2 µL of palladium barcode from the Cell-ID™ 20-Plex Pd Barcoding Kit (Standard BioTools, San Francisco, USA). Cells were then frozen in a freezing solution (90% FBS and 10% DMSO), before shipping to the Flow Cytometry Core Facility at the University of Manchester for data acquisition. Sample analysis was conducted using Cytobank and custom-built, python-based analysis scripts developed in house (**S1B and 1C Figure for QC results**). The antibodies used for labelling were as follow (Standard Biotools, cat No. 201306): Ly6G/C [Gr1] (141^Pr^, clone RB6-8C5, 1/100), CD11c (142^Nd^, clone N418, 1/100), CD69 (145^Nd^, clone H1.2F3, 1/100), CD45 (147^Sm^, clone 30-F11, 1/200), CD11b (148^Nd^, clone M1/70, 1/100), CD19 (149^Sm^, clone 6D5, 1/100), CD3e (152^Sm^, clone 145-2C11, 1/100), TCRβ (169^Tm^, clone H57-597, 1/100), CD44 (171^Yb^, clone IM7, 1/100), CD4 (172^Yb^, clone RM4-5, 1/100), IgM (151^Eu^, clone RMM-1, 1/100), IFNγ (165^Ho^, clone XMG1.2, 1/100).

## Single cell transcriptomics analysis of murine meninges

The single cell suspension obtained from murine meninges after the CD45-PE depletion step was diluted to ∼1,000 cells/μl (in 1X phosphate buffered saline supplemented with 0.04% BSA) and kept on ice until single-cell capture using the 10X Chromium platform. The single cell suspensions were loaded onto independent single channels of a Chromium Controller (10X Genomics) single-cell platform. Briefly, ∼25,000 single cells were loaded for capture using 10X Chromium NextGEM Single cell 3 Reagent kit v3.1 (10X Genomics). Following capture and lysis, complementary DNA was synthesized and amplified (12 cycles) as per the manufacturer’s protocol (10X Genomics). The final library preparation was carried out as recommended by the manufacturer with a total of 14 cycles of amplification. The amplified cDNA was used as input to construct an Illumina sequencing library and sequenced on a Novaseq 6000 sequencers by Glasgow polyomics.

### Read mapping, data processing, and integration

For FASTQ generation and alignments, Illumina basecall files (*.bcl) were converted to FASTQs using bcl2fastq. Gene counts were generated using Cell Ranger v.6.0.0 pipeline against a combined *Mus musculus* (mm10) and *Trypanosoma brucei* (TREU927) transcriptome reference. After alignment, reads were grouped based on barcode sequences and demultiplexed using the Unique Molecular Identifiers (UMIs). The mouse-specific digital expression matrices (DEMs) from all six samples were processed using the R (v4.1.0) package Seurat v4.1.0^38^. Additional packages used for scRNAseq analysis included dplyr v1.0.7, RColorBrewer v1.1.2 (http://colorbrewer.org), ggplot v3.3.5, and sctransform v0.3.3^39^. We initially captured 20,621 cells mapping specifically against the *M. musculus* genome across all conditions and biological replicates, with an average of 30,407 reads/cell and a median of ∼841 genes/cell (**S1 Table and S1D Figure**). The number of UMIs was then counted for each gene in each cell to generate the digital expression matrix (DEM). Low quality cells were identified according to the following criteria and filtered out: *i*) n Feature <200 or >4,000 genes, *ii*) nCounts <200 or >4,000 reads, *iii*) >20% reads mapping to mitochondrial genes, and *iv*) >40% reads mapping to ribosomal genes, v) genes detected < 3 cells. After applying this cut-off, we obtained a total of 19,690 high quality mouse-specific cells with an average of 950 genes/cell (**S1 Table and S1D Figure**). High-quality cells were then normalised using the *SCTransform* function, regressing out for total UMI and genes counts, cell cycle genes, and highly variable genes identified by both Seurat and Scater packages, followed by data integration using *IntegrateData* and *FindIntegrationAnchors*. For this, the number of principal components were chosen using the elbow point in a plot ranking principal components and the percentage of variance explained (30 dimensions) using a total of 5,000 genes, and SCT as normalisation method.

### Cluster analysis, marker gene identification, sub-clustering, and cell-cell interaction analyses

The integrated dataset was then analysed using *RunUMAP* (10 dimensions), followed by *FindNeighbors* (10 dimensions, reduction = “pca”) and *FindClusters* (resolution = 0.7). The resolution was chosen based on in silico analysis using *Clustree*^40^ (**S1E Figure**). With this approach, we identified a total of 19 cell clusters The cluster markers were then found using the *FindAllMarkers* function (logfc.threshold = 0.25, assay = “RNA”). To identify cell identity confidently, we employed a supervised approach. This required the manual inspection of the marker gene list followed by and assignment of cell identity based on the expression of putative marker genes expressed in the unidentified clusters. This was particularly relevant for immune cells detected in our dataset that were not found in the reference atlases used for mapping. A cluster name denoted by a single marker gene indicates that the chosen candidate gene is selectively and robustly expressed by a single cell cluster and is sufficient to define that cluster (e.g., *Cd79a, Cd4, C1qa, Cldn5*, among others). When manually inspecting the gene markers for the final cell types identified in our dataset, we noted the co-occurrence of genes that could discriminate two or more cell types (e.g., DCs, mononuclear phagocytes, fibroblasts). To increase the resolution of our clusters to help resolve potential mixed cell populations embedded within a single cluster and, we subset fibroblasts, DCs, and mononuclear phagocytes and analysed them individually using the same functions described above. In all cases, upon subsetting, the resulting objects were reprocessed using the functions *FindVariableFeatures, RunUMAP, FindNeighbors*, and *FindClusters* with default parameters. The number of dimensions used in each cased varied depending on the cell type being analysed but ranged between 5 and 10 dimensions. Cell type-level differential expression analysis between experimental conditions was conducted using the *FindMarkers* function (*min.pct* = 0.25, *test.use* = Wilcox) and (*DefaultAssay* = “SCT”). For cell-cell interaction analyses, we used CellPhoneDB^41^ and NicheNet^42^ with default parameters using “mouse” as a reference organism, comparing differentially expressed genes between experimental conditions (*condition_oi* = “Infected”, *condition_reference* = “Uninfected”). Pathways analysis for mouse genes were conducted using STRING ^26^ with default parameters. Module scoring were calculated using the *AddModuleScore* function to assign scores to groups of genes of interest (*Ctrl* = 100, *seed* = NULL, *pool* =NULL), and the scores were then represented in feature plots. This tool measures the average expression levels of a set of genes, subtracted by the average expression of randomly selected control genes. The complete gene list used for module scoring derived from previous publications^25^ or from the MatrisomeDB^43^. Once defined, the collated gene list was used to build the module scoring. Raw data and scripts used for data analysis will be made publicly available after peer review.

### Whole mount meningeal preparation and immunofluorescence

After euthanasia, the skull caps were carefully removed using fine tweezers and scissors and placed immediately in 10% neutral buffered Formalin (NFB) for 10 minutes at room temperature. Coronal brain sections were also fixed as above, embedded in paraffin, and processed for Luxol Fast blue used as a proxy to measure the levels of myelin. Following fixation of the skull caps, for immunofluorescence staining, the meninges were detached from the skull caps using a stereotactic microscope and kept at 4°C in 1X PBS containing 0.025% sodium azide until imaging (no longer than one week). For histological analysis, the dura meninges were left attached to the skull and the samples were decalcified prior to embedding in paraffin using neutral EDTA. 2-3 μm skull sections were then prepared for in situ hybridisation experiments or for Masson’s trichrome staining. For immunofluorescence staining, sections were blocked with blocking buffer (1X PBS supplemented with 5% foetal calf serum and 0.2% Tween 20) and incubated with the following primary antibodies at 4°C overnight: REAfinity anti-mouse FITC CD21/35 (Miltenyi, 1:50), rat anti-mouse ER-TR7 (Novus Biologicals, 1:100), REAfinity anti-mouse PE CD3 (Miltenyi, 1:100), REAfinity anti-mouse APC B220 (Miltenyi, 1:100), anti-mouse CD138 PE (BD Bioscience, 1:100). For the detection of ER-TR7, we used an anti-rat antibody coupled with PE (Thermo, 1:500) for 1 hour at room temperature. All the antibodies were diluted in blocking buffer. Slides were mounted with Vectashield mounting medium containing DAPI for nuclear labelling (Vector Laboratories) and were visualized using an Axio Imager 2 (Zeiss). Single molecule fluorescent *in situ* hybridisation (smFISH) experiments were conducted as follow. Briefly, to prepare tissue sections for smFISH, infected animals and naïve controls were anesthetized with isoflurane, decapitated and the skull caps containing the dura mater layer of the meninges were dissected and place on ice-cold 1X HBSS. The skulls were then fixed with 4% paraformaldehyde (PFA) at 4 °C for 15 min, and then dehydrated in 50, 70 and 100% ethanol. After fixation, the skulls caps were decalcified, cut coronally and embedded in paraffin. 5 μm skull cap sections were RNAscope 2.5 Assay (Advanced Cell Diagnostics) was used for all smFISH experiments according to the manufacturer’s protocols. We used RNAscope probes against mouse *Rarres2* on channel 1 (Cat No. 572581), *Cxcl13* on channel 2 (Cat. No. 406311-C2), and Ly6a on channel 3 (Cat. No 427571-C3). All RNAscope smFISH probes were designed and validated by Advanced Cell Diagnostics. For image acquisition, 16-bit laser scanning confocal images were acquired with a 63×/1.4 plan-apochromat objective using an LSM 710 confocal microscope fitted with a 32-channel spectral detector (Carl Zeiss). Lasers of 405nm, 488nm and 633 nm excited all fluorophores simultaneously with corresponding beam splitters of 405nm and 488/561/633nm in the light path. 9.7nm binned images with a pixel size of 0.07um × 0.07um were captured using the 32-channel spectral array in Lambda mode. Single fluorophore reference images were acquired for each fluorophore and the reference spectra were employed to unmix the multiplex images using the Zeiss online fingerprinting mode. All fluorescent images were acquired with minor contrast adjustments where needed, and converted to grayscale, to maintain image consistency.

### Flow cytometry analysis and *ex vivo* stimulation of meningeal-dwelling T cells

To discriminate circulating versus brain-resident immune cells, we performed intravascular staining of peripheral CD45^+^ immune cells, as previously reported^37^. Briefly, a total of 2 μg of anti-CD45-APC-Cy7 antibody (clone 30-F11, in 100 μl of 1X PBS) was injected intravenously ∼3 minutes prior culling. Mice were euthanised as described above and transcardially perfused with ice-cold 0.025% (wt/vol) EDTA in 1X PBS. Whole meninges were enzymatically digested with Collagenase P (1 mg/ml) and DNAse I (1 mg/ml; Sigma) in 1X PBS (HSBB) (Invitrogen) for ∼30 min at 37 °C, according to previously published protocols^44^. Single-cell suspensions were passed through 70 μm nylon mesh filters to remove any cell aggregates. The cell suspension was cleaned up and separated from myelin debris using a Percoll gradient. The resulting fraction was then gently harvested and used as input for *ex vivo* T cell stimulation or used as input for downstream flow cytometry analysis. Briefly, the resulting cell fraction was diluted to a final density of ∼1×10^6^ cells/ml and seeded on a 96 well plate and stimulated with 1X cell Stimulation cocktail containing phorbol 12-myristate 13-acetate (PMA), Ionomycin, and Brefeldin A (eBioSciences^TM^) for 5 hours at 37°C and 5% CO_2_. Upon stimulation, the cells were analysed for the expression of IL-21 and PD-1.

For flow cytometry analysis, meningeal single cell suspensions were resuspended in ice-cold FACS buffer (2 mM EDTA, 5 U/ml DNAse I, 25 mM HEPES and 2.5% Foetal calf serum (FCS) in 1X PBS) and stained for extracellular markers. The list of flow cytometry antibodies used in this study were obtained from Biolegend and are presented in the table below. Samples were run on a flow cytometer LSRFortessa (BD Biosciences) and analysed using FlowJo software version 10 (Treestar). For intracellular staining, single-cell isolates from brain were stimulated as above in Iscove’s modified Dulbecco’s media (supplemented with 1X non-essential amino acids, 50 U/ml penicillin, 50 μg/ml streptomycin, 50 μM β-mercaptoethanol, 1 mM sodium pyruvate and 10% FBS. Gibco). Cells were then permeabilized with a Foxp3/Transcription Factor Staining Buffer Set (eBioscience) and stained for 30 min at 4 °C. The anti-mouse GP38 (1:100) and the LTβ (monoclonal antibody BBF6^45^; 10 μg/ml) antibodies were kindly provided by Dr. Lalit Kumar Dubey (QMUL). For the detection of LTβ in CD4^+^ T cells, we used a goat anti-hamster (Armenian) IgG coupled to FITC as secondary antibody (Biolegend; 1:200). For the detection of GP38 we used a Syrian hamster-anti mouse GP38 followed by anti-Syrian hamster secondary antibody coupled to APC/alexa647 (Jackson ImmunoResearch; 1:100). We used the following commercially available antibodies from Biolegend: CD45-APC-Cy7 (clone 30-F11, 2 μg/100 μl 1X PBS i.v.), TER-119-APC-Cy7 (clone TER-119; 1/400), CD19-APC-Cy7 (clone 1D3/CD19; 1/400), F4/80-APC-Cy7 (clone BM8; 1/400), F4/80-PE Dazzle 594 (clone BM8; 1/400), CD3-APC (clone 17A2; 1/400), CD4-FITC (clone GK1.5; 1/400), PD1-BV711 (clone 29F.1A12; 1/400), CXCR5-BV421 (clone L138D7; 1/200), IL-21-PE (clone 3A3-N2; 1/200), CD45-BV711 (clone 30-F11; 1/400), CD31-BV421 (clone 8.1.1; 1/100), CD21/35-PE Dazzle 594 (clone 7E9; 1/100), MAdCAM-1-Alexa Fluor 488 (clone MECA-367; 1/100), CD19-Alexa Fluor 488 (clone 6D5; 1/400), CD138-PE (clone 281-2; 1/200), IgG-BV421 (clone Poly4053; 1/200), IgM-BV711 (clone RMM-1; 1/200), CD8a-BV421 (clone QA17A07; 1/400), I-A/I-E-PerCP-Cy5.5 (clone M5/114.15.2; 1/400)

## ELISPOT assays

ELISPOT tests to measure *ex vivo* the frequency of meningeal antibody secreting cells (ASCs) was performed using the ELISpot Flex IgM– and IgG-HRP (Mabtech) as followed. After generating single cell suspensions from meningeal preparations, a total of 50,000 cells per well were seeded on 96-wells multiscreen-HA filter plates (Millipore) coated with 50 μg/ml of either whole *T. brucei*, prepared in house, or mouse brain lysate (Novus Biologicals) to determine the presence of *T. brucei*– and mouse brain-reactive antibody secreting cells, respectively. Wells coated with 50 μg/ml BSA (Sigma) were also included as negative controls. After seeding the cells, the plates were incubated for 16 hours at 37°C, and 5%CO_2_ covered in foil to avoid evaporation. In parallel, plates coated with 15 μg/ml affinity-purified goat anti-mouse IgM and IgG were also analysed in parallel to measure the frequency of total IgM and IgG ASCs. For this, we used a total of 25,000 cells per well and incubated as before. Spots were enumerated with an Immunospot analyser (CTL, Germany).

## Detection of autoreactive IgM and IgG by ELISA

Serum samples from naïve and infected animals at 30 days post-infection were used to examine the presence of mouse brain lysate-specific IgM and IgG using a colorimetric approach. For this purpose, polysorb ELISA plates (Biolegend) were coated overnight with 50 μg/ml either *T. brucei* Antat 1.1E whole cell lysate prepared in house, or mouse brain lysate (Novus Biologicals) in 1X coating buffer (Biolegend). After extensive washes with 1X ELISA washing buffer (Biolegend), total mouse IgM or IgG were detected in mouse serum (1:50 to 1:10,000 dilution in 1X PBS) or human CSF (1:400 in 1X PBS) by using Horseradish peroxidase-conjugated antibodies specific for mouse IgM (Thermo) or IgG (all isotypes; Sigma) using the recommended concentrations, and the resulting absorbance was detected at 450 nm using an ELISA Multiskan plate reader (Thermo).

### Detection of host antigens detected by autoantigens

Blood samples were collected by cardiac puncture from naïve mice (*n* = 3 mice) or at 30dpi (*n* = 3 mice) were and place on EDTA tubes, from which serum was obtained. In parallel, we screened CSF samples collected from 1^st^ stage sleeping sickness patients (*n* = 3 patients) and 2^nd^ stage patients (*n* = 4 patients). Due to ethical constraints, we did not have access to CSF samples from African healthy donors. Therefore, we obtained CSF samples from healthy Caucasian donors (*n* = 2 donors) from the University of Edinburgh Brain and Tissue Bank. Autoantibodies were assessed using a commercial microarray-based platform (GeneCopoeia). Briefly, mouse serum or human CSF samples were hybridised to distinct microarray spots containing 120 native host and viral antigens spotted onto nitrocellulose fibers (adhered to glass slides). Next, the slides were incubated with fluorescently-coupled anti-IgG or anti-IgM secondary antibodies, and microarrays were scanned using a GenePix 4400A microarray scanner. Raw fluorescence data was normalized to PBS controls on each slide. The data presented in the heatmaps are normalised signal-to-noise ratios.

## Data availability

The transcriptome data generated in this study have been deposited in the Gene Expression Omnibus (GSE229436; Reviewer’s token: mzitckmgbtmtxqz). The processed transcript count data and cell metadata generated in this study, as well as the code for analysis, are available at Zenodo (https://doi.org/10.5281/zenodo.7814657). Additional data and files can also be sourced via Supplementary Tables.

## Code availability

The processed transcript count data and cell metadata generated in this study, as well as the code for analysis, are available at Zenodo (DOI: 10.5281/zenodo.7814657).

## Statistical analysis

All statistical analyses were performed using Graph Prism Version 8.0 for Windows or macOS, GraphPad Software (La Jolla California USA). The data distribution was determined by normality testing using the Shapiro-Wilks test. Where indicated, data were analysed by unpaired Student’s t-test, Mann-Whitney test, or one-way analysis of variance (ANOVA). Data were considered to be significant where *p* <0.05. For the *in vivo* experiments, we matched sex and age of the mice in experimental batches using a block design including randomisation of experimental units. Data collection and analysis were not performed blindly to the conditions of the experiment due to the specific requirements of the UK Home Office project licence.

## Results

### The murine meninges are colonised by a diversity of immune cells during chronic *T. brucei* infection

We and others have shown an increase in meningeal infiltration and meningitis during the chronic stage (25dpi onwards) in experimental infections with *T. brucei*^10,13^. Previous studies have shown that mouse meninges are colonised by CD2^+^ T cells and CD11c^+^ dendritic cells (DCs) during chronic *T. brucei* infection^9^, although a catalogue of the immune interactions spanning beyond these compartments is lacking. To fill this gap in knowledge, we used an integrative multi-omics approach that combined CyTOF and 10X Chromium single cell transcriptomics (**Figure 1A**) to understand the complexity of the immune interactions taking place in the meningeal space during chronic *T. brucei* infection. In addition to unbiasedly cataloguing the cells involved in this process, this approach also allowed us to identify transcriptional pathways involved in the anti-parasitic responses in the meninges with as much resolution as possible. We focussed on characterising the dura mater as this has been previously shown to contain the vast majority of the meningeal CD45^+^ immune cells^42^, as well as parasites during the chronic stages of the infection^9^. To ensure we captured the diversity of resident immune cells and meningeal stroma with as much confidence as possible, we selectively removed all circulating CD45^+^ immune cells using a magnetic sorting approach (**Figure 1A and S1A Figure**). In brief, we labelled circulating CD45^+^ cells by intravenous injection with an anti-CD45^+^ antibody coupled to phycoerythrin (PE). All the circulating CD45^+^ were then isolated using an anti-PE antibody coupled to magnetic beads. After extensive perfusion, the remaining circulating (PE^+^ cells) cells were removed leaving behind resident CD45^+^ cells (PE^-^ cells), as well as stromal cells, including fibroblasts and cells associated with the vasculature and the lymphatic system.

**Figure 1.**
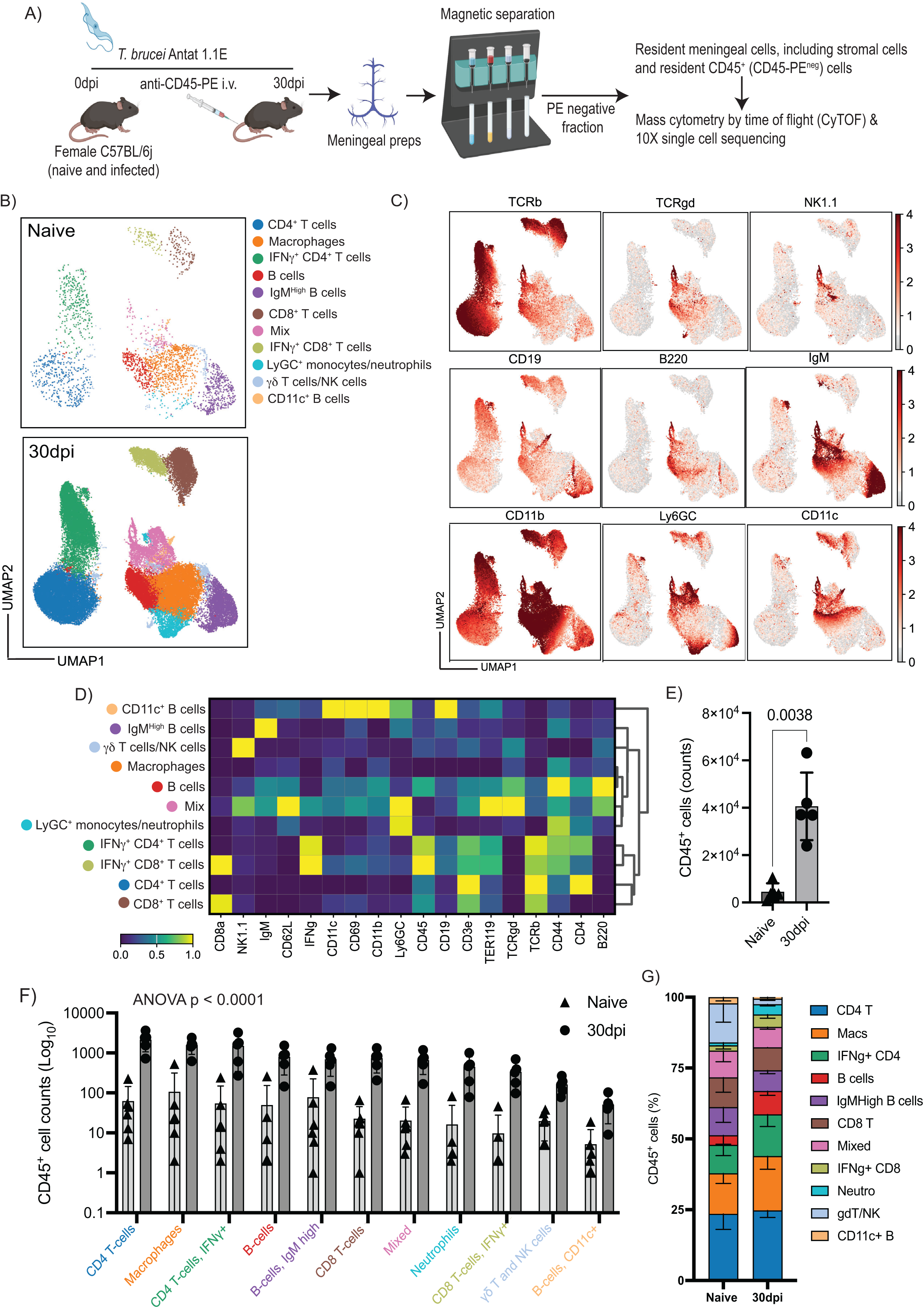
CyTOF confirms the expansion of innate and adaptive immune cells in the murine meninges during chronic *T. brucei* infection. A) Overview of the experimental approach applied in this work. An anti-CD45-PE antibody was injected *i.v.* prior to cull, followed by magnetic sorting using anti-PE antibodies to obtain a fraction of resident meningeal cells that were used as input for single cell transcriptomics using the 10X Chromium platform. **B)** Uniform manifold approximation and projection (UMAP) visualisation of the CyTOF immunophenotyping in murine meninges from naïve and animals at 30dpi. The UMAP plot represents pooled data from n = 5-6 mice/group. **C)** Normalised protein expression level of markers used to define T cells and NK cells (TCRβ, TCRγ8, NK1.1), B cells (CD19, B220, IgM), and myeloid cells (CD11b, Ly6G/C, and CD11c). **E)** Quantification of CD45^+^ cells in the murine meninges by CyTOF at 30dpi (*n* = 5 mice/group). Parametric two-sided *T* student test: a *p* value < 0.05 was considered significant. **D)** Unsupervised cell annotation from CyTOF data using a combination of several marker genes. The expression level is normalised to the average of the expression within the group. **F)** Quantification of the different populations of immune cells in the naïve and infected murine meninges (*n* = 5-6 mice/group). A parametric ANOVA test with multiple comparison was used to estimate statistically significant pairwise comparisons. A *p* value < 0.05 is considered significant. Data in all panels are expressed as mean ± SD. Data points indicate biological replicates for each panel. **G)** Bar chart depicting the frequency of the different immune populations identified in the murine meninges by CyTOF.

Firstly, we decided to explore the broad immunological landscape in the meninges of mice at 30dpi, when infection in the CNS is well established^9–13^. For this reason and given the scarcity of cells typically obtained from the meninges, CyTOF was used as this approach enables detection of a wide range of cell types. Our CyTOF data was composed of T cells (CD4^+^ and CD8^+^ T cells, including IFNγ^+^ subsets, NK and γο T cells), B cells (including IgM^High^ B cells and CD11c^+^ B cells), and myeloid cells including neutrophils and macrophages (**Figure 1B-D**, **S1B and S1C Figure**), consistent with previous work^2^. Overall, we detected a significant increase in the number of CD45^+^ cells (**Figure 1E**), and an increased in number of the various immune subsets in the murine meninges at 30dpi compared to naïve controls (**Figure 1F**), without noticeable changes in cell frequency **(Figure 1G**). Together, these data suggested the expansion and/or recruitment of resident immune cells into the meninges in response to infection.

To gain an understanding of the transcriptional responses triggered in the meninges in response to infection, and to identify potential interactions between various meningeal cells during infection, we performed single cell RNA sequencing of meningeal preparations from mice at 30 dpi (*n* = 2 pools; 2 mice/pool) and naive controls (*n* = 2 pools; 2 mice/pool). Using this approach and after removing cells catalogued as low quality (**Materials and Methods**), we obtained a total of 19,690 high quality cells, from which 1,834 cells derived from naïve meninges and 17,856 cells from infected meninges, with an average of 605 genes per cell from naïve samples and 1,297 genes per cell from infected samples (**Figure 2A**, **Figure S1D and S1E, and Materials and Methods**). As expected, our single cell meningeal atlas encompasses stromal and immune cells, most of which have previously been reported in the murine meninges^2,46^. Within the stromal compartment, we identified two populations of *Col1a1^+^* fibroblasts, *Ccl19^+^ Rarres2^+^* mural cells, and *Wvf^+^ Pecam1^+^* endothelial cells (**Figure 2A and 2B**). Within the immune compartment, we identified five populations of mononuclear phagocytes (MNPs 1 to 5), characterised by the expression of putative myeloid cell markers genes such as *Ccl8, C1qa, Aif1, Adgre1* and *Cd14*, amongst others, as well as conventional DCs (cDCs; *Xcr1, Zbtb46, Clec9a, Flt3* and *Itgae*) (**Figure 2A and 2B**). Additionally, we also detected T cells (*Trac, Cd3g, Cd3e, Cd4, Icos, Cd8a, Gzmb, Gzmk*), granulocytes, including *S100a8^+^ Ngp^+^ Cd177^+^*neutrophils, and two populations of B cells (*Cd79a, Cd79b*) that expressed markers of canonical plasma cell markers (e.g., *Sdc1*) (**Figure 2A and 2B**). Lastly, we detected a small proportion of cells (<0.5%) with high expression levels of haemoglobin (*Hba-a1, Hbb-bt*) and genes typically related to neurons (*Neurod1, Neurod4*) (**Figure 2A**, **2B, and S2 Table**). Notably, we observed a robust increase in the frequency of most of the cells within the immune compartment (**Figure 2C**) consistent with the CyTOF dataset. Together, these analyses are consistent with profound alterations in the cellular makeup of the murine meninges during chronic *T. brucei* infection.

**Figure 2.**
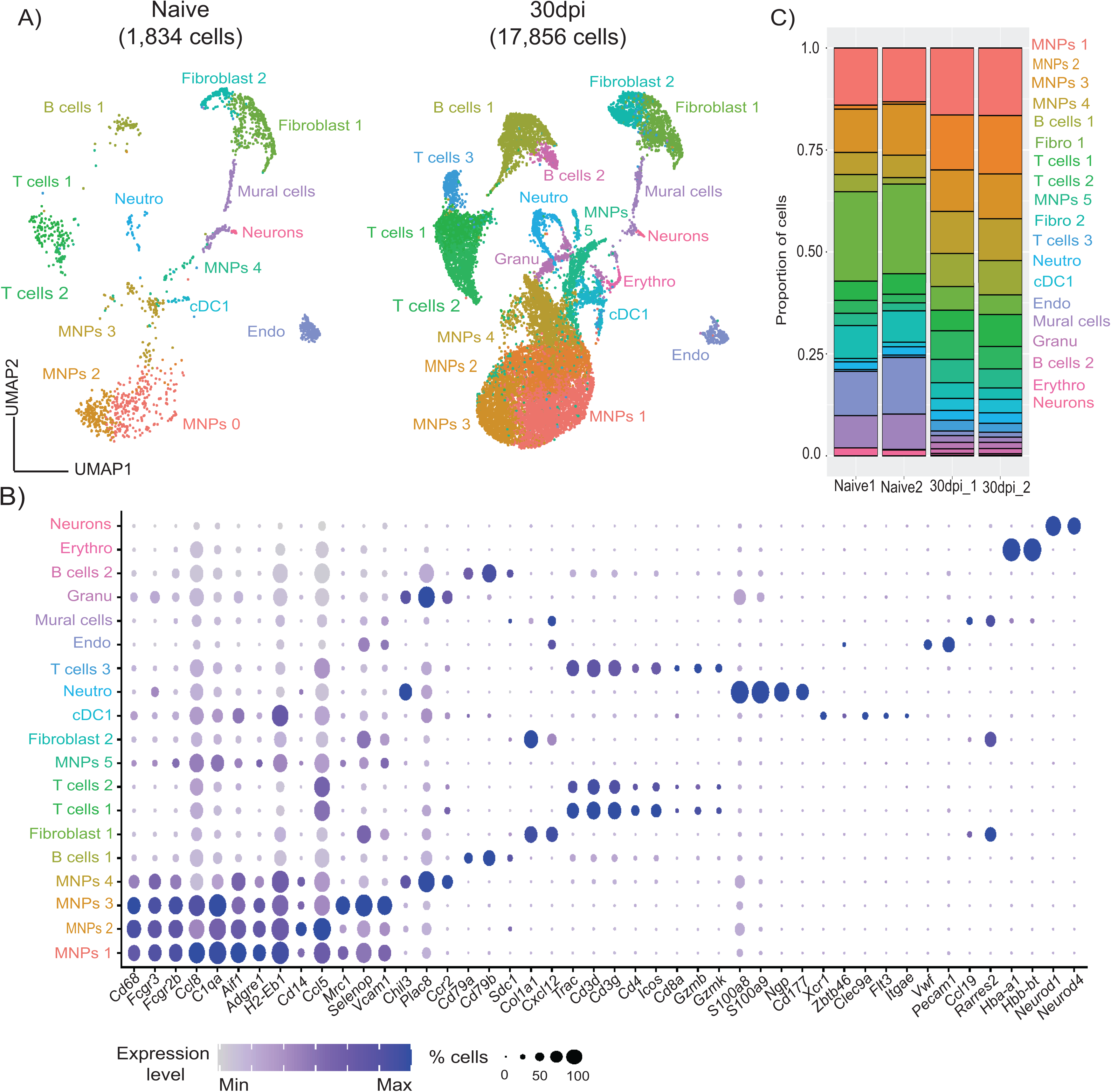
Single cell atlas of the murine meninges during chronic *T. brucei* infection. A) A total of 19,690 high-quality cells were used for dimensionality reduction, resulting in uniform manifold approximation and projection (UMAP) for the single cell transcriptome profiling from naïve (n = 2 pools; 1,834 high quality cells in total) and infected meninges (n = 2 pools; 17,856 high quality cells in total). **B)** Dotplot representing the expression levels of top marker genes used to catalogue the diversity of cell types in our single cell dataset. **C)** Frequency of the different cell types detected in the murine meninges analysed in this study.

## Meningeal *Rarres2^+^ Ly6a*^+^ fibroblasts acquire lymphoid tissue stroma-like properties during chronic *T. brucei* infection

Next, we focussed on the meningeal fibroblasts. The meningeal fibroblasts are a heterogeneous cell population encompassing transcriptionally, spatially, and potentially functionally distinct units critical for meningeal immunity^47–50^, but their responses to chronic protozoan infections remains to be elucidated. Therefore, we first asked whether the three fibroblast clusters (Fibroblast 1, Fibroblast 2, and mural cells) identified in figure 2 contained discreet clusters of cell populations that were not resolved by the top-level clustering. After sub-clustering, we identified 2,088 cells (547 and 1,541 cells from naïve and infected meningeal preparations, respectively), distributed across eight discrete subsets (**Figure 3A**). These clusters expressed marker genes putatively associated with fibroblasts from the dura mater, including *Mgp, Gja1, Fxyd5 and Col18a1* (**Figure 3B**)^48^, but low or undetectable levels of markers proposed to be associated with arachnoid mater fibroblasts (e.g, *Clnd11, Tbx18, Tagln*) or pia mater (e.g., *Lama2, S100a6, Ngfr*) (**Figure 3B**)^48^. These observations suggested that the majority of the fibroblasts within our dataset were likely to be derived from the dura mater layer of the meninges^51^. These fibroblast clusters also expressed *Col1a1, Col1a2, Pdgfra* and *Pdgfrb* to various degrees but lacked *Pecam1* (encoding for CD31), suggesting that these cells are of mesenchymal origin rather than endothelial^50,51^ (**Figure 3C**). Furthermore, cells within clusters 0 to 5 expressed *Rarres2*, suggesting that these are meningeal pericyte-like fibroblasts^50^, in addition to *Ly6a* (which encodes for the Stem Cell antigen-1, *Sca1*) (**Figure 3C**), suggesting they are likely to retain progenitor properties^50^. We thus classified these cells as *Ly6a^+^*fibroblasts (**Figure 3A to 3C**). Clusters 0 to 3 also expressed *Pdpn*, which encodes for GP38, and *Col6a2*, which is recognised by the antibody ER-TR7^52^, and were thus defined as fibroblast reticular cells (FRCs)-like *Ly6a^+^* fibroblasts 1 to 4 (**Figure 3C**). Cells within cluster 4 expressed high levels of the antioxidant protein Fth1 in addition to *Rarres2* and *Ly6a* and were labelled as *Fth1^+^ Ly6a^+^* fibroblasts (**Figure 3C**). Cells within cluster 5 also expressed *Ccl19* and *Aldh1a2,* recently shown to be associated with lymphoid stroma in the milky spots^53^, in addition to *Ly6a*, and were thus classified as *Aldh1a2^+^ Ccl19^+^* fibroblasts (**Figure 3C**). Cells within cluster 6 expressed high levels of *Acta2* (which encodes for α-smooth muscle actin), as well as *Tnn, Postn*, and *Mmp13*, and were thus classified as myofibroblasts^50^. Lastly, cells within cluster 7 expressed *Slc38a2* and *Slc47a1*, consistent with the phenotype of dura and leptomeningeal fibroblasts^54,55,56^ recently reported to be enriched in several transporters, and were thus assigned as *Ly6a^-^* dura fibroblasts (**Figure 3C**).

**Figure 3.**
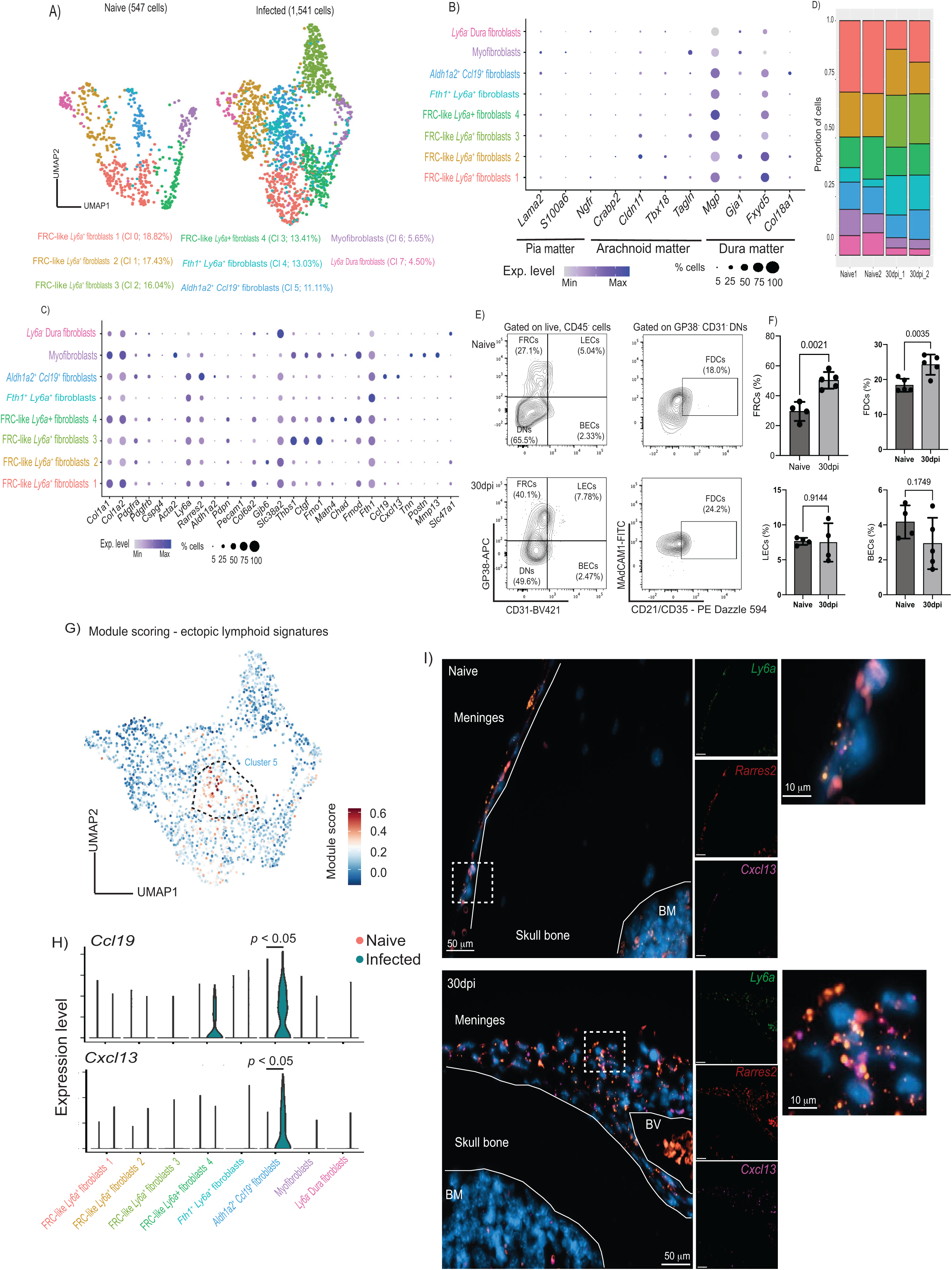
The meningeal stromal compartment acquires lymphoid-like properties and provide cues for immune cell recruitment during chronic *T. brucei* infection A) Uniform manifold approximation and projection (UMAP) of 2,088 high-quality cells within the fibroblast clusters were re-analysed to identify a total eight subclusters. **B)** Dot plot representing the expression levels of marker genes previously reported to be enriched in fibroblasts from different meningeal layers, including dura mater, pia mater, and arachnoid mater. **C)** Dot Plot representing the expression levels of top marker genes used to catalogue the diversity of cell types in our single cell dataset. **D)** Frequency plot the different fibroblast clusters detected in the murine meninges in naïve and infected samples (*n* = 2 pools per experimental condition). **E)** Representative flow cytometry analysis of meningeal stromal cells from naïve and infected meningeal preparations (left panel), and the corresponding quantification **(F)**. A The data in all panels is expressed as mean ± SD. Data points indicate biological replicates for each panel. A parametric T test was employed to assess significance between experimental groups. A *p* value < 0.05 is considered significant. FRC, fibroblast reticular cells; LECs, lymphatic endothelial cells; BECs, blood endothelial cells; FDCs, follicular dendritic cells. **G)** Module scoring for genes typically associated with lymphoid tissues. **H)** Violin plot depicting the expression of two chemokines associated with lymphoid tissues such as *Ccl19* and *Cxcl13*. **I)** Single molecule fluorescent *in situ* hybridisation (smFISH) demonstrating the presence of *Ly6a^+^ Rarres2*^+^ cells that express *Cxcl13* in the dura mater of the meninges of infected mice. DAPI is included as a nuclear marker. The skull bone, the bone marrow (BM), and the meninges are indicated. Scale bar, 50 μm.

We also found heterogeneous responses to infection within the meningeal fibroblast subset, including cells that display features of FRC and B cell zone reticular cells (BCRs) in response to infection. Cells within cluster 0, 1, 2, 3, and to a lesser extent cells within cluster 5 upregulated genes associated with FRCs^57^ in response to infection, including *Pdpn, Pdgfra, Pdgrfb, Vim*, and *Col6a3*, as well as secreted factors such as *Vefga* and *Tnfsf13b* (encoding for BAFF) (**S3A Figure**). Cells within clusters 3, 5, and 6 upregulated genes associated with BRCs^58^ during infection, including *Cxcl13* (Cluster 5), *Itga7, Ltbr* (cluster 5 and 6)*, Madcam1* (cluster 5), *Cr2* (cluster 3 and 6), and *Tnfsf11* (encoding for RANKL) in cluster 3 (**Figure 3C and S3B Figure**). Based on the expression pattern observed across the fibroblast subset in response to infection, we catalogued clusters 0, 1, 2, and 3 as FRC-like cells, and cells within clusters 5 and 6 as BRC-like cells (**S3C Figure**). Lastly, cells within cluster 2, described as a mature population of pericytes, were exclusively detected in the infected meninges and were characterised by the expression of genes associated with blood vessel development (*Fgfr, Thbs1, Fgf2*), as well as leukocyte chemotaxis and myeloid differentiation (*Cxcl19*) (**Figure 3C and S3 Table**), suggesting *de novo* expansion in response to infection. Additionally, all of the FRC-like pericytes are predicted to be involved in extracellular matrix (ECM) remodelling in the meninges, including collagen and proteoglycan deposition, as well as secretion of factors involved in ECM production, as demonstrated by module scoring analysis using the MatrisomeDB database^43^ (**Figure S3D Figure**), which encompasses a curated proteomic dataset of ECM derived from a wide range of murine tissues. Consistent with these *in silico* predictions, we observed a consistent pattern of fibroblastic reactions and collage deposition in the dura meninges from infected mice compared to naïve controls (**S3E Figure**), further indicating an extensive meningeal ECM remodelling triggered in response to infection.

Our results so far indicate that the dura mater layer of the meninges contains a diverse population of stromal cells, including GP38^+^ and GP38^-^ stromal cells that resemble the stroma of other lymphoid tissues and ECM remodelling. Consistent with our scRNAseq data, using flow cytometry, we observed a significant expansion of GP38^+^ FRCs, and a distinctive population of MAdCAM1^+^ CD21/CD35^+^ cells indicative of the presence of FDC-like cells in the infected murine meninges compared to naïve controls (**Figure 3E-F**; **Gating strategy in 2A Figure**), without significant changes in the lymphatic endothelial cells (LECs) or blood endothelial cells (BECs) (**Figure 3E-F**). Furthermore, using module scoring analysis, which allows us to assess global gene signatures associated with a gene set or pathway (in this case, ectopic lymphoid tissue signatures), we were able to identify that cells within cluster 5 were enriched for genes associated with FDC-like function and stromal lymphoid tissues, including *Ccl19 and Cxcl13*, compared to the other fibroblast clusters (**Figure 3H**), and may be derived from *Ly6a^+^* pericytes with a precursor capacity as previously reported^59,60^. Indeed, using *in situ* hybridisation on independent tissue sections, we were able to confirm the presence of *Ly6a^+^ Rarres2^+^* cells that expressed *Cxcl13^+^* during infection (**Figure 3I**). Together, our data demonstrate the presence of a rich diverse fibroblast population, encompassing *Ly6a^+^ Rarres2^+^* FRC-like pericytes, including *Aldh1a2^+^ Ccl19^+^* FRC-like pericytes, myofibroblasts, *Fth1^+^* fibroblasts, and perivascular dura fibroblasts. Our data also suggests that chronic *T. brucei* infection induces an extensive remodelling of the meningeal stroma compartment, resulting in the expansion of FRCs and FDC-like cells without significant changes in the vasculature (LECs and BECs).

## Meningeal mononuclear phagocytes are predicted to be involved in antigenic presentation and chemotaxis during chronic *T. brucei* infection

The cells within the myeloid compartment in the murine meninges act as a critical first line of defence against insults and were clearly expanded during the chronic stage of the infection (**Figure 1 and 2**). To resolve the mononuclear phagocyte (MNPs) compartment in more detail, we analysed these populations individually. In total, we obtained 10,760 cells organised into five major clusters: cluster 0 (33.8%), cluster 1 (25.04%), cluster 2 (19.7%), cluster 3 (13.4%), and cluster 4 (8.10%) (**Figure 4A**). Since clusters 0 and 2 expressed high levels of *Mrc1* (encoding for CD206) and the anti-inflammatory molecule *Il18bp*, and *Siglec1* (encoding for CD169), we labelled these clusters as *Cd206*^+^ border-associated macrophages (BAMs) (**Figure 4B and S4 Table**). Clusters 1 and 3 contained the immune sensors *Cd14* and *Tlr2*, in addition to *Mertk, Adgre1*, and *Ly6c2*. We therefore labelled these clusters as monocyte-derived macrophages (MDMs). Lastly, cells within cluster 4 expressed high levels of mitochondrial-associated transcripts (e.g., *mt-Co1, mt-Co2, mt-Atp6*), in addition *Itgal*, *Sirpa, Cd274, Nfkbia, Sell*, and *Cd44* (**Figure 4B and S4 Table**), and were labelled as metabolically active mononuclear phagocytes (maMNPs) (**Figure 4A and B**). We also observed that *Cd206*^+^ BAMs and cells within the MDMs 1 cluster expressed high levels of *H2-Aa, Sirpa, Csf1r, Cxcl16,* and *Adgre1,* which encodes for F4/80^61^ (**Figure 4B and S4 Table**). Under homeostatic conditions, the murine meninges were dominated by *Cd206*^+^ BAMs, in agreement with previous reports^62^ (**Figure 4C**). However, during infection, there was a significant expansion of MDMs (**Figure 4C**), suggesting that the murine meninges were populated by circulating monocytes during chronic *T. brucei* infection, consistent with previous reports^10^. Cell-cell interaction analyses predicted that meningeal MNPs establish significant interactions with other cell types, including T cells, *via* antigenic presentation (**Figure 4D to 4F**), likely driving T cell activation locally, as previously proposed^2,10^. Consistent with our *in silico* predictions, based on the expression level of F4/80, we identified two populations of CD11b^+^ myeloid cells, that we defined as CD11b^+^ F4/80^high^ (resembling *Cd206*^+^ BAMs and MDMs 1) and CD11b^+^ F4/80^low^ (resembling MDMs 2 and maMNPs) (**Figure 4B**, **4G, and 4H**). During infection, there was a significant increase in the frequency of CD11b^+^ F4/80^low^ MNPs, whereas the CD11b^+^ F4/80^high^ MNPs population decreased in frequency (**Figure 4G and 4H; Gating strategy in S2B Figure**). However, in both cases, we noted a significant increase in the expression of MHC-II in both CD11b^+^ F4/80^high^ and CD11b^+^ F4/80^low^ MNPs (**Figure 4G and 4H**). Together, our results indicate that the resident population of meningeal myeloid cells expand upon infection (e.g., either as a result of local myeloid proliferation or *via* the recruitment of monocytes to the meningeal space) likely driving both cell recruitment *via* chemotaxis and antigen presentation to CD4^+^ T cells. Our data are consistent with and complementary to independent reports focusing on the ontogeny and dynamics of BAMs and MNPs under homeostasis^2^ and during the onset and resolution of *T. brucei* infection^10^.

**Figure 4.**
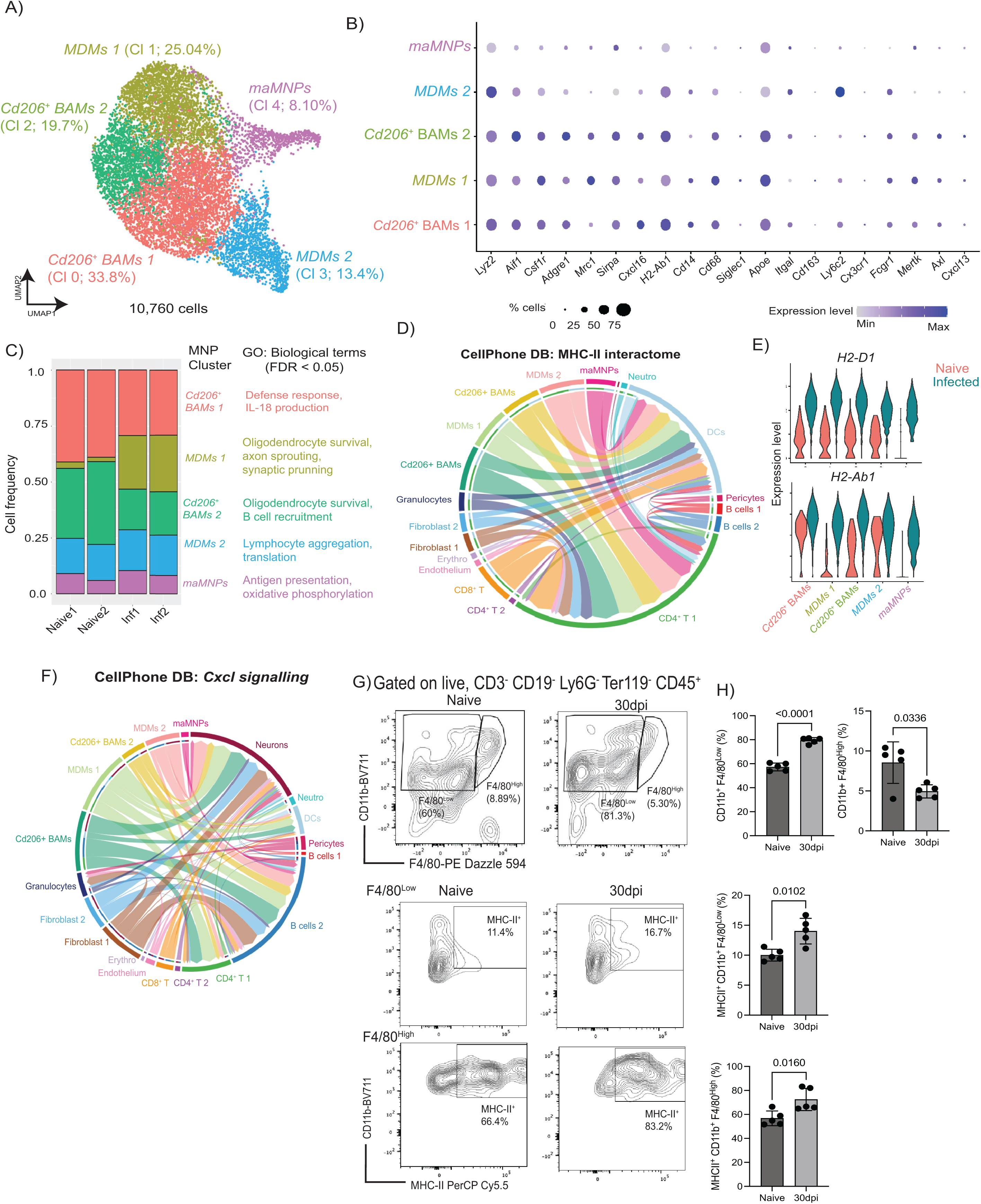
**Heterogeneity of meningeal mononuclear phagocytes during chronic T. brucei infection**. **A)** Uniform manifold approximation and projection (UMAP) of 10,760 high-quality mononuclear phagocytes (MNPs) from naïve (n = 54 cells) and infected meninges (n = 10,706 cells). **B)** Expression level of top genes defining different populations of meningeal MNPs. **C)** Frequency plot depicting the relative abundance of the five MNPs subclusters identified in the murine meninges during *T. brucei* infection. **D)** Cell-cell interaction network via MCH-II signalling axis. **E)** Violin plot depicting the expression level of *H2-D1* and *H2-Ab1*, two of the most upregulated MHC-II associated genes within the myeloid compartment. **F)** Cell-cell interaction network *via Cxcl* signalling axis. **G)** Representative flow cytometry analysis and quantification (**H**) of CD11b^+^ F4/80^High^ and F4/80^Low^ myeloid cell populations, as well as MHC-II^+^ myeloid cells, in the murine meninges in response to *T. brucei* infection. Data in all panels are representative from two independent experiments and is expressed as mean ± SD (*n* = 5 mice/experimental group). Data points indicate biological replicates for each panel. A parametric T test was employed to assess significance between experimental groups. A *p* value < 0.05 was considered significant.

## The murine meninges contain T_FH_-like cells during chronic *T. brucei* infection

The accumulation of inflammatory T cell subsets in the meninges has been reported in CNS infections with *T. brucei*^9^, but their features and effector functions remain unresolved. Our top level single cell analysis identified three discreet T cell clusters based on the expression of *Trac, Cd3e, Cd3g, Cd4*, and *Cd8a* (**Figure 2A**). To resolve the meningeal T cell compartment in more detail, we re-clustered the T cells and repeated the dimensionality reduction analysis. Within the resident meningeal T cell compartment, we identified four main transcriptional clusters, characterised by the expression of *Trbc1, Cd4* (cluster 0, 1, and 2), and *Cd8a* (cluster 3) (**Figure 5A and 5B, and and S5 Table**). Several of the genes observed in the CD4^+^ T cells were putatively associated with a T_FH_ like phenotype, including *Icos, Pdcd1* (encoding for PD-1), *Cxcr4, Ctla4, Maf*, *Nr4a1, Csf1*, *Tox2, Cxcr5, Bcl6*, as well as the cytokines *Ifng* and *Il21* (**Figure 5B and 5C**). To confirm this, we first examined the presence of T_FH_-like T cells in the meninges *in vivo* using flow cytometry. Consistent with our *in silico* predictions, we detected a significant increase in the frequency of resident CXCR5^+^ PD1^+^ CD4^+^ T cells in the murine meninges in response to chronic *T. brucei* infection compared to naïve controls (**Figure 5D and 5E; Gating strategy in S2C Figure**). Furthermore, *ex vivo* stimulation assays demonstrated that chronic *T. brucei* infection results in a significant expansion of meningeal PD1^+^ CD4^+^ T that express IL-21 compared to naïve controls (**Figure 5F and 5G**), further indicating their T_FH_-like phenotype.

**Figure 5.**
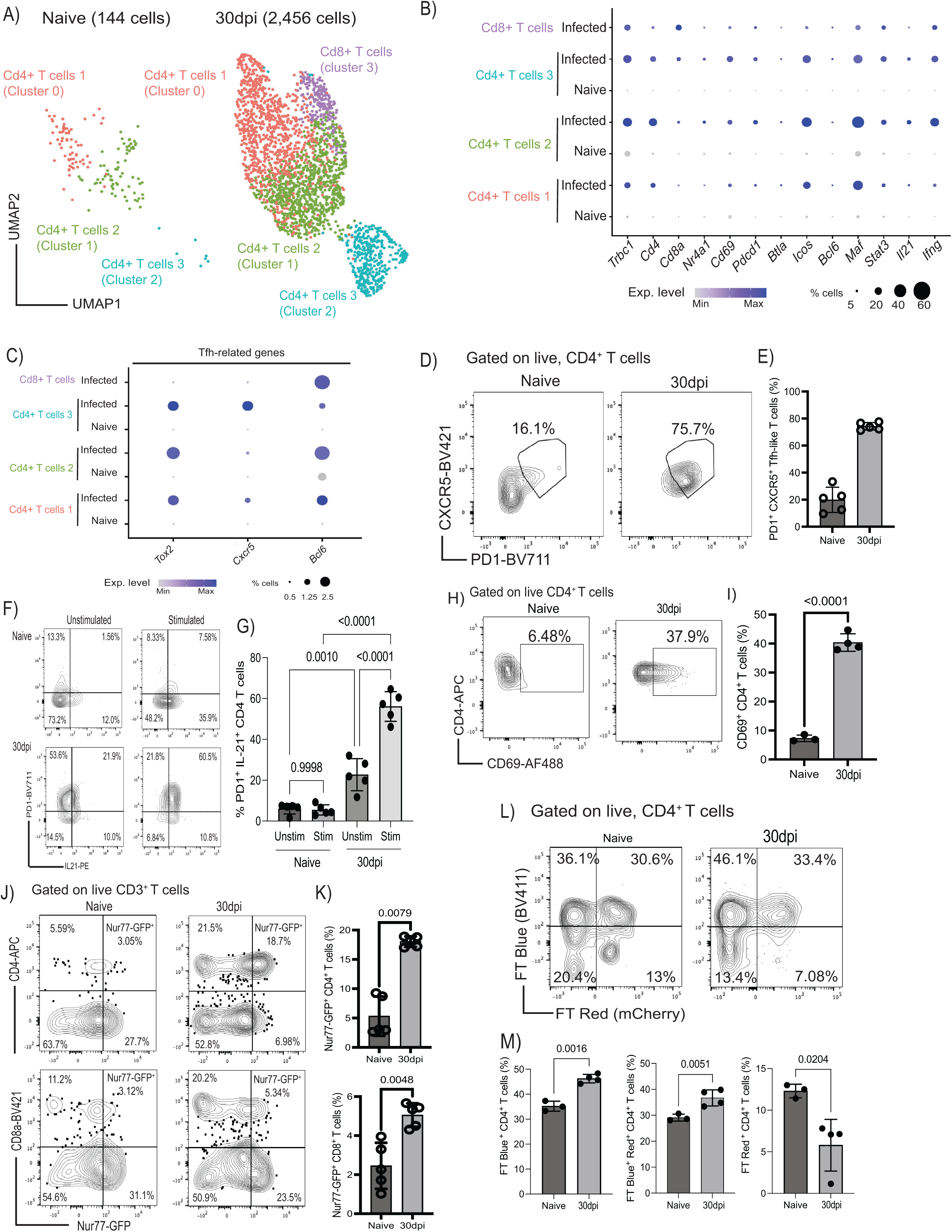
**Accumulation of PD1^+^ CXCR5^+^ T**_FH_**-like CD4^+^ T cells in the meninges during chronic *T. brucei* infection A)** Uniform manifold approximation and projection (UMAP) of 1,742 high-quality T cells from naïve (n = 147 cells) and infected meninges (n = 2,691 cells). **B)** Dot plot depicting the expression level of marker genes for all the T cell subsets identified in (**A**). The dot size corresponds to the proportion of cells expressing the marker genes, whereas the colour indicates the level of expression. **C)** as in (B) but depicting marker genes associated with T_FH_ cells. The dot size corresponds to the proportion of cells expressing the marker genes, whereas the colour indicates the level of expression. **D)** Representative flow cytometry analysis and quantification (**E**) the presence of CXCR5^+^ PD1^+^ CD4^+^ T cells in the murine meninges from naïve and 30dpi (*n* = 5 mice/group). Data points indicate biological replicates for each panel. A parametric T test was employed to assess significance between experimental groups. A *p* value < 0.05 is considered significant. **F)** *Ex vivo* T cell activation from naïve and infected murine meninges to measure the expression of PD-1 and IL-21. Unstimulated controls are also included. **G)** Quantification of the flow cytometry data from the *ex vivo* stimulation assay in (F). Data points indicate biological replicates for each panel and are representative from two independent experiments. A parametric T test was employed to assess significance between experimental groups. A *p* value < 0.05 was considered significant. **H)** Representative flow cytometry analysis and quantification **(I)** of the frequency of CD69^+^ CD4^+^ T cells in the murine meninges in response to infection. Data points indicate biological replicates for each panel and is representative from two independent experiments. A parametric T test was employed to assess significance between experimental groups. A *p* value < 0.05 was considered significant. **J)** Representative flow cytometry analysis to determine TCR engagement in CD4^+^ (top panel) and CD8^+^ (bottom panel) T cells *in situ* in the meninges during chronic infection with *T. brucei* using the Nur77^GFP^ reporter mouse line. **K)** Quantification of the flow cytometry data from **(J)**. A *p* value < 0.05 is considered significant. Data points indicate biological replicates for each panel and are representative from two independent experiments. A parametric T test was employed to assess significance between experimental groups. A *p* value < 0.05 is considered significant. **L)** Representative flow cytometry analysis to determine TCR engagement in CD4^+^ (top panel) in the Nur77^Tempo^ reporter mouse line. In this model, T cell activation dynamics can be discriminated between *de novo* (FT blue^+^) versus historical (FT red^+^) MHC-dependent TCR engagement. **M)** Quantification of the flow cytometry data from **(L)**. Data points indicate biological replicates for each panel and are representative from two independent experiments. A parametric T test was employed to assess significance between experimental groups. A *p* value < 0.05 was considered significant.

Our data so far also indicate that meningeal ecosystems promote T cell activation *via* antigen presentation (**Figure 4D**). Indeed, both our *in silico* prediction and flow cytometry experiments demonstrated an expansion of CD69^+^ CD4^+^ T cells in the infected meninges compared to naïve controls (**Figure 5B**, **5H, and 5I**), strongly suggesting local activation. To examine whether meningeal T cells were activated *in situ* during infection, we initially utilised Nur77^GFP^ reporter mice^63^. In this model, GFP expression is used as a proxy for MHC-dependent TCR engagement resulting in T cell activation^63^. We observed a significant increase in the frequency of *Nur77*-GFP^+^ CD4^+^ and CD8^+^ T cells (**Figure 5J**, **5K; Gating strategy in S2C Figure**), indicating local T cell activation within the murine meninges. To further resolve whether T cell activation occurs *in situ*, we used the newly reported *Nur77*^Tempo^ mice, a novel murine reporter line in which the expression of a fluorescent timer (FT) protein is driven by *Nur77* expression^64,65^. This model enabled the discrimination of newly activated (FT blue^+^), persistent (FT blue^+^ red^+^), and arrested (FT red^+^) T cells based on MHC-dependent TCR engagement^64,65^. We observed a significant increase in the frequency of newly activated and persistent CD4^+^ T cells, and a concomitant reduction in the frequency of arrested CD4^+^ T cells in the meninges in response to infection compared to naïve controls (**Figure 5L and 5M**), indicating that most CD4^+^ T cells are actively partaking in the local immune response, likely via antigenic presentation. We also observed a significant increase in the frequency of newly activated meningeal CD8^+^ T cells, but a reduction in both persistent and arrested CD8^+^ T cells, perhaps indicating that the CD8^+^ T cell responses are transitory (**S4A and S4B Figure**). This pattern of local T cell activation was also detected in the CD69^+^ CD4^+^ T cells, in which we detected a higher frequency of newly activated CD69^+^ CD4^+^ T cells and less of arrested CD69^+^ CD4^+^ T cells (**S4C and S4D Figure),** altogether indicating the meningeal T cells are newly activated in the meninges *in situ*. These observations are consistent with previous studies showing that CD4^+^ T cells actively patrol the meningeal landscape^2^. Taken together, these results demonstrate that the meningeal CD4^+^ T cell population undergoes newly and persistent MHC-dependent TCR engagement in the meninges, promoting local responses *in situ*, but the CD8^+^ T cell responses seem more transitory. These responses are likely to provide all the necessary signals for activation, likely resulting in T cell differentiation towards the observed a T_FH_-like phenotype during chronic *T. brucei* infection.

## The murine meninges contain plasmablasts/plasma cells and GL7^+^ CD95^+^ GC-like B cells during chronic infection

Previous studies have demonstrated that B cells represent a major immune population in the meninges^42,46^, although their dynamics during chronic infection are not yet understood. We previously observed the expression of *Cxcl12* in dura and arachnoid meningeal fibroblasts, which is critical for the differentiation and survival of early B cells in the bone marrow^66,67^. Thus, we next explored the diversity of B cells in our dataset. The majority of meningeal B cells detected in our dataset derived from the infected meninges (1,688 cells out of 1,742 total B cells) (**Figure 2**). These cells expressed high levels of genes associated with plasmablasts and plasma cells such as such as *Jchain*, *Prdm1* (which encodes for BLIMP-1)*, Sdc1* (encoding CD138), *Ighm*, and *Irf4* (**Figure 6A**). Flow cytometry experiments further confirmed that the vast majority of the meningeal B cells correspond to plasmablasts and plasma cells, and to a lesser extent CD19^+^ B cells (**Figure 6B; Gating strategy in S2D Figure**). Some of the marker genes identified within the B cell clusters, such as *Pcna, Mki67, Ub2c and Ighg2* and *Ighg3*, are typically associated with cell replication and class-switched B cells (**S2 Table**). These genes are critical for affinity maturation and class switching during GC reactions^68^. Because the transcriptional signatures observed within the B cell clusters were consistent with the presence of extrafollicular GC-like reactions, we next examined this at the protein level. We first exploited the Nur77^GFP^ reporter mouse line to measure BCR engagement within the meninges. Given that Nur77^GFP^ expression in GC B cells is proposed to be markedly reduced compared to activated B cells *in vivo*^69^, and that *Nur77* restrains B cell clonal dominance during GC reactions^70^, this reporter line can be used to examine extrafollicular GC-like reactions. In line with these studies, we detected significantly fewer GFP^+^ CD19^+^ B cells during infection compared to naïve controls (**Figure 6C and 6D, and S2D Figure**), implying that upon infection, the meningeal B cells undergo GC-like reactions. Intriguingly, the meningeal B cells also expressed *Cd38* and *Fas* (**Figure 6A**), similar to dark zone (GFP^low^) GC B cells^69^. Consistent with these observations, we detected a significant accumulation of GL7^+^ CD95/Fas^+^ cells within the CD19+ B cell compartment (**Figure 6E and 6F**), further corroborating the presence of GC-like B cells in the murine meninges. Consistent with the GC-like and the transcriptional profile, we observed an increase in the frequency of IgG^+^ CD19^+^ B cells in the murine meninges at 30dpi compared to naïve controls (**Figure 6G and 6H**). In the spatial context, we observed clusters of CD3^+^ T cells, B220^+^ B cells and CD21/35^+^ FDCs in the murine meninges that were not readily detectable in naïve animals (**Figure 6I and S5 Figure**), suggesting the presence of immunological aggregates similar to those observed in tertiary lymphoid tissues^26,27^. Together, our data indicates the presence of class-switched plasma cells/plasmablasts as well as GC-like CD19^+^ B cells in close proximity to CD3^+^ T cells and CD21^+^/CD35^+^ FDC-like cells in the murine meninges in response to *T. brucei* infection.

**Figure 6.**
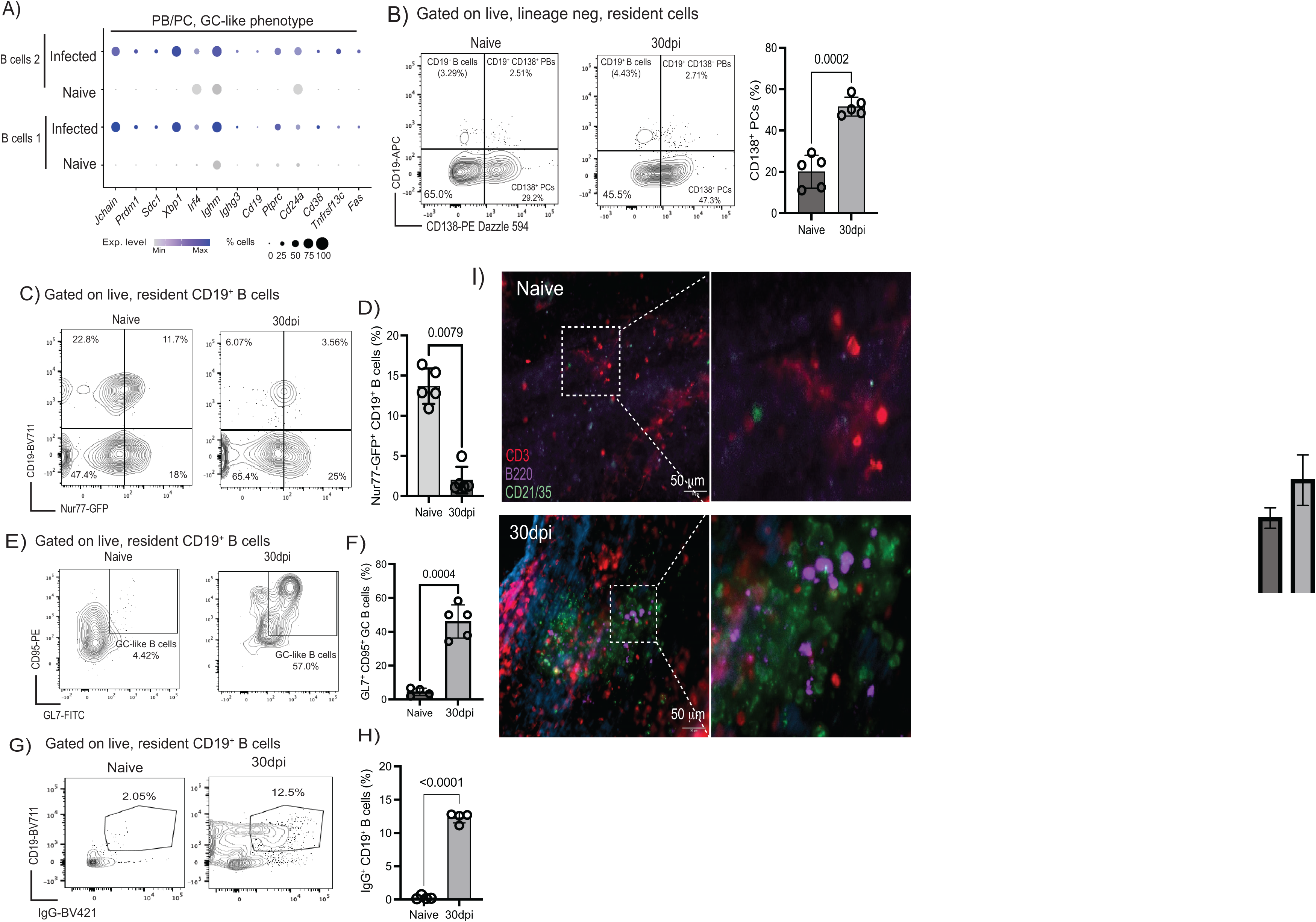
**The murine meninges contain class-switched B cells in proximity to FDC-like cells**. **A)** Dotplot representing the expression level of top marker genes for the meningeal B cells, including *bona fide* markers of plasma cells (*Jchain, Prdm1, Sdc1, Xbp1, Ighm)* as well as activation and GC-like phenotype (*Cd24a, Cd38, Tnfrsf13c, Fas*). The dot size corresponds to the proportion of cells expressing the marker genes, whereas the colour indicates the level of expression. **B) Left panel;** Representative flow cytometry analysis (left panel) and quantification (right panel) to measure meningeal B cells (CD19^+^ CD138^-^ cells), plasmablasts (CD19^+^ CD138^+^), and plasma cells (CD19^-^ CD138^+^). **Right panel;** quantification of flow cytometry data showing the expansion of meningeal CD138^+^ plasma cells in response to infection (*n* = 5 mice/group). Data points indicate biological replicates for each panel and are representative from two independent experiments. A parametric T test was employed to assess significance between experimental groups. A *p* value < 0.05 was considered significant. **C)** Representative flow cytometry analysis to determine BCR engagement on meningeal B cells *in situ* during chronic infection with *T. brucei* using the *Nur77*^GFP^ reporter mouse line. **D)** Quantification of flow cytometry data showing the reduction in the frequency of *Nur77*^GFP+^ CD19^+^ B cells in the meninges in response to infection (*n* = 5 mice/group). Data points indicate biological replicates for each panel and are representative from two independent experiments. A parametric T test was employed to assess significance between experimental groups. A *p* value < 0.05 was considered significant. **E)** Representative flow cytometry analysis to determine the presence of GC-like B cells in the murine meninges based on the co-expression of GL7 and CD95. **D)** Quantification of GL7^+^ CD95^+^ GC-like CD19^+^ B cells in the meninges in response to infection (*n* = 5 mice/group). Data points indicate biological replicates for each panel and are representative from two independent experiments. A parametric T test was employed to assess significance between experimental groups. A *p* value < 0.05 was considered significant. **G)** Representative flow cytometry analysis to determine the presence of IgG^+^ CD19^+^ B cells in the murine meninges in response to infection. **H)** Quantification of GL7^+^ CD95^+^ GC-like CD19^+^ B cells in the meninges in response to infection (*n* = 4 mice/group). Data points indicate biological replicates for each panel and are representative from two independent experiments. A parametric T test was employed to assess significance between experimental groups. A *p* value < 0.05 was considered significant. **I)** Representative imaging analysis of whole-mount meninges from naïve (left) and infected (right) of CD21/CD35^+^ follicular dendritic cells (green), as well as CD3d^+^ T cells (red) and B220^+^ B cells (purple). DAPI was included as nuclear staining. Scale = 50 μm.

### *T. brucei* infection results in the accumulation of meningeal autoreactive B cells

The accumulation of meningeal B cells has been reported in several autoimmune disorders such as neuropsychiatric lupus^25^ and multiple sclerosis^28^ where they are responsible for the generation of autoantibodies that are linked to the pathology associated with these disorders. However, it is unclear whether chronic *T. brucei* infection also results in the accumulation of autoreactive B cells in the meningeal spaces. We reasoned that in addition to generating B cell clones able to generate antibodies specific to *T. brucei*, these local GC-like reactions taking place within the meningeal space might also result in the development of autoreactive B cells. To directly test this hypothesis, we examined the presence of meningeal resident IgG^+^ antibody secreting cells (ASCs) able to recognise *T. brucei* and mouse brain lysates using ELISpot. We observed a significant accumulation of total IgG^+^ ASCs in the murine meninges (**Figure 7A and 7B**), consistent with our flow cytometry data (**Figure 6G-H**). We also detected a significant accumulation of IgG^+^ ASCs able to recognise *T. brucei* and mouse brain but not BSA (**Figure 7A and 7B**), indicative of the presence of autoreactive ASCs in the murine meninges during infection. Interestingly, splenocytes from animals at 30dpi or naïve controls did not contain autoreactive IgG^+^ ASCs (**S6A Figure**), suggesting that the mouse brain-specific autoreactive ASCs may arise locally within the meninges or within the CNS environment. Histological analysis of the corresponding murine brain sections revealed extensive IgG deposition in the infected brain compared to naïve controls, in particular in the leptomeninges and the cortex (**Figure 7C**). The IgG antibody deposition observed in the cerebral cortex in response to chronic infection was accompanied by a significant demyelination, particularly in the cerebral cortex, internal capsule, and thalamic tracts (**Figure 7D**, **and S6B and S6C Figure**). Additionally, we detected the presence of high IgM and IgG titres in the serum of infected animals able to react to mouse brain antigens compared to naïve controls (**Figure 7E**), further corroborating our histological and ELISpot findings. It is important to note that the binding of circulating IgG antibodies to the murine brain does not seem to be restricted to areas with high parasite accumulation (e.g., lateral ventricles) (**Figure 7F**). In humans, in the cerebrospinal fluid of 2^nd^ stage *gambiense* HAT patients from North Uganda we observed significant levels of autoreactive IgM and IgG antibodies able to recognise human brain lysates, but not BSA (**Figure 7G**, **and S6 Table**), consistent with our findings in experimental infections. Taken together, our data suggest that meningeal B cells undergo affinity maturation locally within the meninges or the CNS space to generate IgG^+^ ASCs directed against both *T. brucei* and the mouse brain (and in gambiense HAT patients), is associated with cortical and white matter demyelination, and results in autoimmunity.

**Figure 7.**
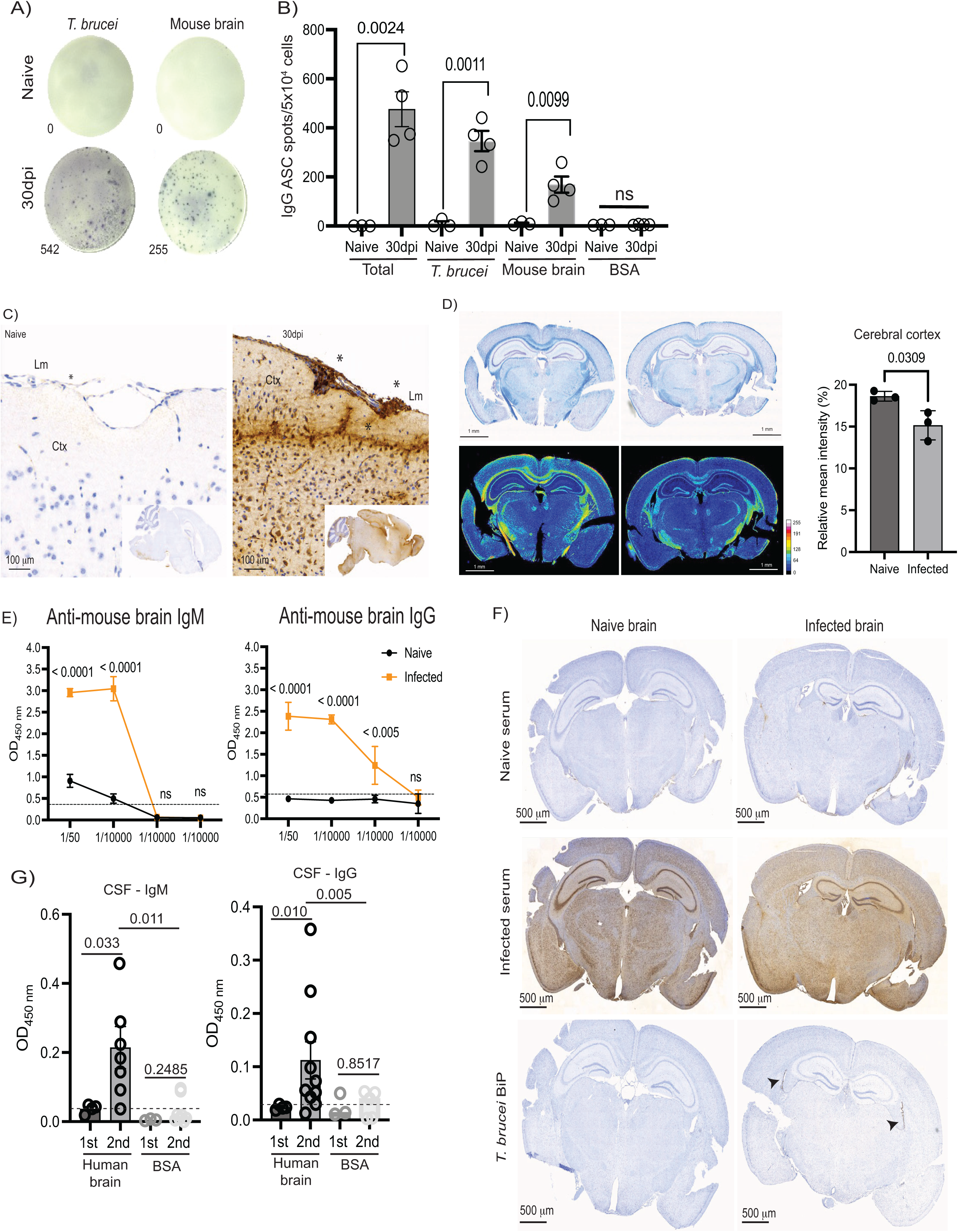
**The murine meninges contain autoreactive IgG^+^ antibody secreting cells (ASCs) during chronic *T. brucei* infection**. **A)** Representative ELISpot images depicting mouse brain-specific IgG^+^ ASCs from naïve and infected murine meninges after 30dpi with *T. brucei*. **B)** Quantification of ELISpot results including total IgG^+^, *T. brucei*-specific IgG^+^, and mouse brain-specific IgG^+^ antibody secreting cells. Wells coated with BSA were included as negative controls. A p value < 0.05 was considered significant. **C)** Immunohistochemistry analysis to determine IgG^+^ deposition in the mouse brain from naïve (left) and infected (right) mouse brain sagittal sections. The sections were stained with an anti-mouse IgG antibody coupled to HRP to measure the overall distribution of IgG in the brain. The asterisks denote areas of intense IgG deposition in the leptomeningeal space, as well as in the upper layers of the cerebral cortex, exclusively detected in the infected brain. Lm, leptomeninges; Ctx, cerebral cortex. **D) Left panel:** Representative Luxol fast blue (LFB) staining from naïve (left) and infected (right) animals at 30dpi as a proxy to measure myelin. Lower panels show the tissue heatmap of the mean pixel intensity. **Right panel:** Percentage of demyelination, calculated here as a reduction in the average of the relative LFB intensity, was calculated from three independent experiments (n = 3-4 mice/repeat). A parametric T test was employed to assess significance between experimental groups. A *p* value < 0.05 was considered significant. **E)** Serum titters of mouse brain-specific IgM and IgG antibodies in naïve and infected samples as measured by ELISA. The dotted line represents the average of the optical density detected in naïve controls. A *p* value < 0.05 was considered significant. **F)** Immunohistochemistry analysis to determine the presence of circulating brain autoreactive IgG antibodies in serum from naïve (top panels) and infected animals (middle panels) in the mouse brain from naïve (left) and infected (right) mouse coronal brain sections. Staining with the *T. brucei*-specific protein BiP (bottom panels) is also included to highlight accumulation of parasites in the lateral ventricles (arrowheads). **G)** ELISA analysis of human brain-autoreactive IgM and IgG antibodies in cerebrospinal fluid (CSF) from sleeping sickness patients from 1^st^ stage and 2^nd^ stage HAT (CSF dilution 1:400). Wells coated with BSA (5μg/ml) were included as controls. A *p* value < 0.05 was considered significant.

## LTβ receptor signalling controls the accumulation of meningeal FDCs and autoreactive B cells during chronic *T. brucei* infection

LTβ receptor (LTβR) signalling is critical for the formation, induction, and maintenance of lymphoid tissues under homeostasis and disease^71–73^. This process requires interactions between the LTα_1_β_2_ heterodimer and its cognate receptor LTβR to induce broad effects on FDC maintenance, promoting a favourable microenvironment promoting GC reactions on B cells^71–73^. Furthermore, expression of LTα in the meninges causes *de novo* ectopic lymphoid tissue formation and neurodegeneration in a model of myelin oligodendrocyte glycoprotein-induced experimental autoimmune encephalitis^74^. Our data so far indicate that the murine meninges develop ectopic lymphoid aggregates that display many features of LTβ-driven lymphoid tissue formation, including the presence of FDCs like structures, T_FH_ T cells, and GC-like B cells with evidence of somatic hypermutation. Thus, we hypothesised that LTβR-signalling plays a similar role in the formation of meningeal lymphoid aggregates and coordinating the meningeal responses to chronic *T. brucei* infection. The gene which encodes the LTβR, *Ltbr,* was expressed myeloid cells, endothelial cells, granulocytes, and fibroblasts in the meninges (**Figure 8A**), indicating that LTβR signalling may occur at multiple levels within the murine meninges. Similarly, LTβ (encoded by *Ltb*) is primarily expressed by the CD4^+^ T cell clusters and to a lesser extent by cDCs, neutrophils, CD8^+^ T cells, and B cells (**Figure 8A**). Using flow cytometry, we detected a significant increase in the frequency of CD4^+^ T cells expressing LTβ (**Figure 8B and 8C**), consistent with their T_FH_ phenotype^27^. Next, we investigated the role of LTβR-signalling in the maintenance of local immunological responses within the meningeal stroma. For this, mice were treated prior and during *T. brucei* infection with a LTβR-Ig fusion protein to prevent the interaction of the ligands LTα_1_β_2_ and LIGHT (encoded by *Tnfsf14*) with LTβR (**Figure 8D**)^75^. LTβR-Ig treatment resulted in mice unable to control systemic parasitaemia as efficiently as mice treated with an irrelevant antibody or untreated mice (**S7A Figure**), and in a worsening in the clinical scoring (**S7B Figure**), mirroring previous work using *Ltb*^-/-^ mice infected with *T. brucei* in the context of intradermal infections^76^. Furthermore, LTβR-Ig treatment significantly impaired the expansion of meningeal FDCs (**Figure 8E and 8F**), and a significant accumulation of meningeal LECs compared to naïve controls, which can be attributed to changes in frequencies within other stromal compartments (**Figure 8E and 8F**). Using ELISpot, we observed that LTβR-Ig treatment significantly impaired the expansion of both IgM^+^ (**Figure 8G and S7C-E Figure**) and IgG^+^ ASCs, including *T. brucei*– and mouse brain-specific ASCs (**Figure 8G**), consistent with a central role for LTβR-signalling in the formation of B cell follicles and GCs within secondary lymphoid organs and ectopic lymphoid tissues^59,60,71,76^. Lastly, LTβR-Ig treatment significantly impaired the formation of perivascular FDC-B cell clusters (**Figure 8H**), consistent with previous reports^59,60^, and prevented the cortical demyelination typically observed in response to chronic infection (**Figure 8I and S7E Figure**). Together, these data demonstrate that LTβR signalling is required for stromal responses and B cell accumulation and maturation in the meninges during infection with *T. brucei*, further highlighting that the meninges depend on classical lymphoid tissue-associated signalling pathways to coordinate local immune responses to infections. Furthermore, the fact that LTβR-Ig treatment rescued the cortical demyelination observed in response to infection suggests that the meningeal ectopic lymphoid aggregates are indeed pathogenic.

**Figure 8.**
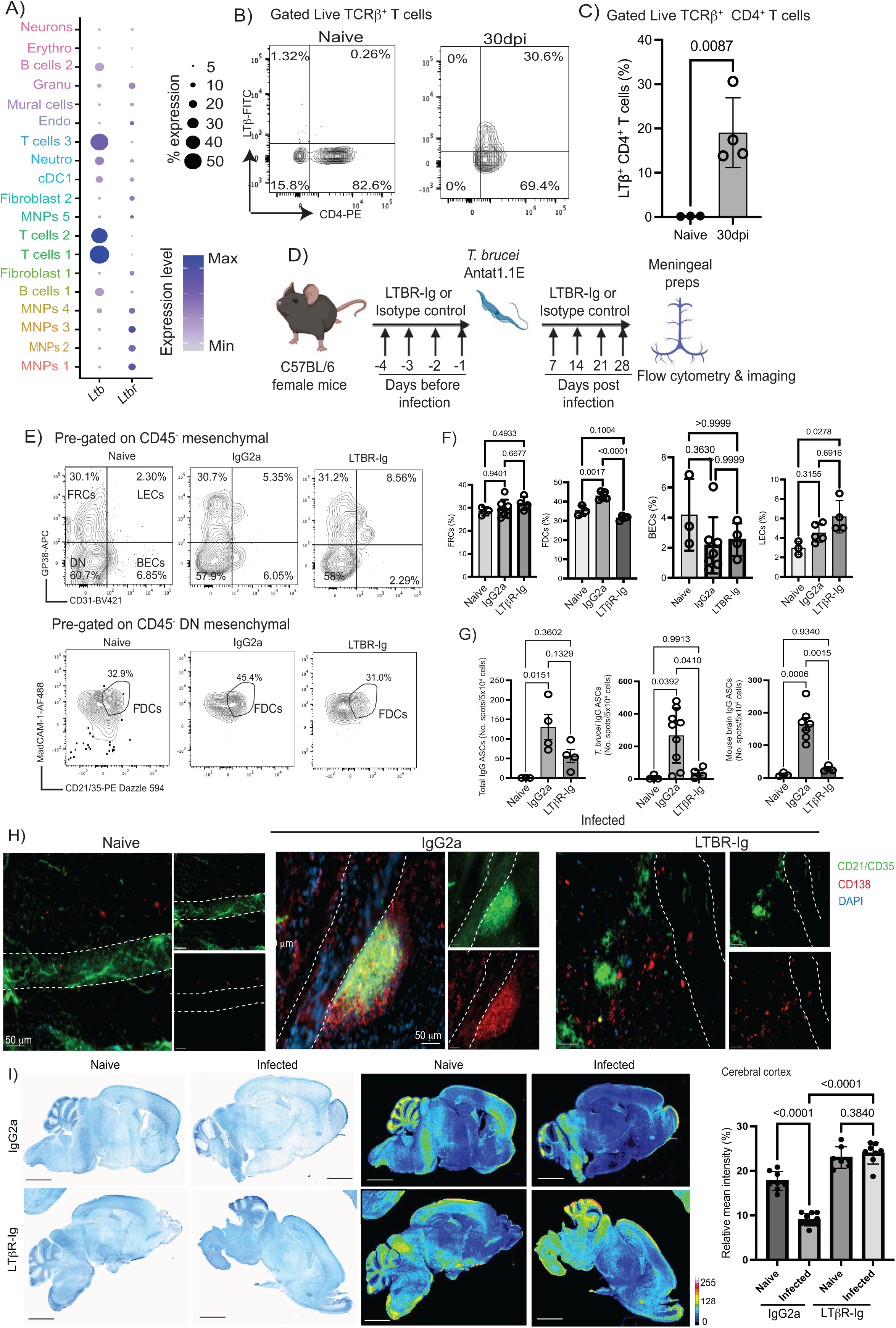
LTβ receptor signalling is critical to sustain FDC-like networks and autoreactive B cells in the murine meninges in response to *T. brucei* infection. **A)** Dot plot depicting the expression level of *Ltb*, and its cognate receptors *Traf2* and *Ltbr*. The dot size corresponds to the proportion of cells expressing the marker genes, whereas the colour indicates the level of expression. **B)** Representative flow cytometry analysis depicting the expression of LTβ in CD4^+^ T cells in naïve and mice chronically infected with *T. brucei* (30dpi). **C)** Quantification of flow cytometry analysis (n = 4 mice/group). A *p* value < 0.05 is considered significant. **D)** Overview of the experimental approach applied to block LTβR signalling *in vivo* during chronic *T. brucei* infection. **E)** Representative flow cytometry analysis of the murine meningeal stroma in naïve and mice chronically infected with *T. brucei* (30dpi). FDC-like cells were gated from the double CD45^-^ mesenchymal cells. **F)** Quantification of the different components of the stroma in naïve and infected meningeal preparations (n = 4 mice/group). A *p* value < 0.05 was considered significant. FRC, fibroblast reticular cell; LEC, lymphatic endothelial cell; BEC, blood endothelial cell; DN, double negative. **G)** Quantification of ELISpot results including total IgG^+^ (left panel), *T. brucei*-specific IgG^+^ (middle panel), and mouse brain-specific IgG^+^ antibody secreting cells (right panel) in naïve mice, mice treated with an irrelevant IgG2a antibody, and mice treated with LTBR-Ig (*n* = 4 – 9 mice/group). A *p* value < 0.05 was considered significant. **H)** Representative immunofluorescence analysis of whole mount meningeal preparation labelling CD138^+^ plasma cells (red) and CD21/CD35^+^ FDC-like cells (green) in naïve or at 30 days post-infection. Each of the fluorescent channels are shown individually, and DAPI was included to detect cell nuclei. Scale bar = 50 μm. **I) Left panel:** Representative Luxol fast blue (LFB) staining from naïve (left) and infected (right) animals at 30dpi as a proxy to measure myelin. **Middle panel:** The tissue heatmap of the mean pixel intensity is also shown. **Right panel:** Percentage of demyelination, calculated here as a reduction in the mean grey intensity of the LFB staining, was calculated from two independent experiments (n = 4-5 mice/experiment). A parametric ANOVA with multiple comparisons was employed to assess significance between experimental groups. A *p* value < 0.05 was considered significant.

## Infection-induced autoantibodies recognise a broad range of host antigens, including myelin basic protein

Given that our data so far indicate that chronic *T. brucei* infection results in the generation of autoantibodies, we next decided to examine the nature of antigens recognised by these autoantibodies. To achieve this, we employed a targeted array of 120 antigens known to be identified in autoimmune disorders, from systemic lupus erythematous to multiple sclerosis. Our data indicates that circulating IgG autoantibodies found in infected samples significantly recognised a total of 18 antigens (15% of the antigen array), including structural proteins (e.g., collagen VI, vitronectin, nucleolin, Histone H3), cytokines (e.g., GM-CSF), components of the complement system (e.g., C3, C1q), intracellular antigens (e.g., ssDNA, ssRNA, mitochondrial antigen), and most importantly myelin basic protein (MBP) (**Figure 9A**, **S7 Table, and S8A Figure**). To further understand whether the same pattern of autoreactive antibodies is observed in sleeping sickness patients, we screened CSF samples collected from patients during the 1^st^ (haemolymphatic) stage and 2^nd^ (meningoencephalitic) stage. As observed in mice, our results highlighted a broad range of host antigens recognised by IgG autoantibodies in the CSF exclusively detected during the 2^nd^ stage of the disease (**Figure 9B**, **S7 Table, and S8B Figure**). More specifically, we detected reactivity against 51 antigens (42.5% of the antigen array), including several structural proteins, cytokines (e.g., TGFβ1, TNFα, IL-12, TPO), intracellular antigens (e.g., histones, nucleosome-related proteins, mitochondrial antigen), structural proteins (e.g., collagens, vitronectin), amongst others (**Figure 9B**). Interestingly, as observed in mice, we also detected the presence of autoantigens able to bind host proteins associated with either parasite control or pathology, such as proteins of the complement system (e.g., C1q, C3a) and nervous system-associated proteins (e.g., MBP and muscarinic receptor) (**Figure 9B**). Indeed, a total of 8 antigens (13.1% of the antigen array) were commonly identified by autoreactive antibodies in infected mouse serum and human CSF from 2^nd^ stage sleeping sickness patients, which are known to be diagnostic markers of autoimmune disorders such as systemic lupus erythematosus, Sjogren’s syndrome, scleroderma, rheumatoid arthritis, and multiple sclerosis^77–79^ (**Figure 9C**). To further validate our findings, we examined the presence of circulating antibodies against MBP in an independent cohort of sleeping sickness patients from DRC that included both patients with an active infection (“cases”) and samples obtained from patients post-treatment (“treated) (**Figure 9D**). Consistent with the data obtained from CSF biopsies, we observed that sleeping sickness patients with an active infection have significantly higher titres of serum IgG against MBP compared to healthy African controls (**Figure 9D**). Interestingly, most of the samples obtained from patients post-treatment display basal levels of anti-MBP antibody titres and show no significant differences with healthy African controls, suggesting that treatment with anti-parasitic chemotherapy prevents the accumulation of anti-MBP autoantibodies in humans. However, we noted that 30% of the treated patients maintained higher titres of anti-MBP antibodies in circulation, which might be due to: a) failure to effectively clear parasites post-treatment and thus considered to be relapsing cases; b) the presence of memory B cells that sustain anti-MBP autoantibody secretion; c) or a combination of both. Taken together, our mid-throughput targeted screening identified a myriad of host antigens recognised by infection-induced autoantibodies in both chronically infected mice and 2^nd^ stage sleeping sickness patients, potentially indicative of a complex autoimmune disorder affecting several organs including the CNS.

**Figure 9.**
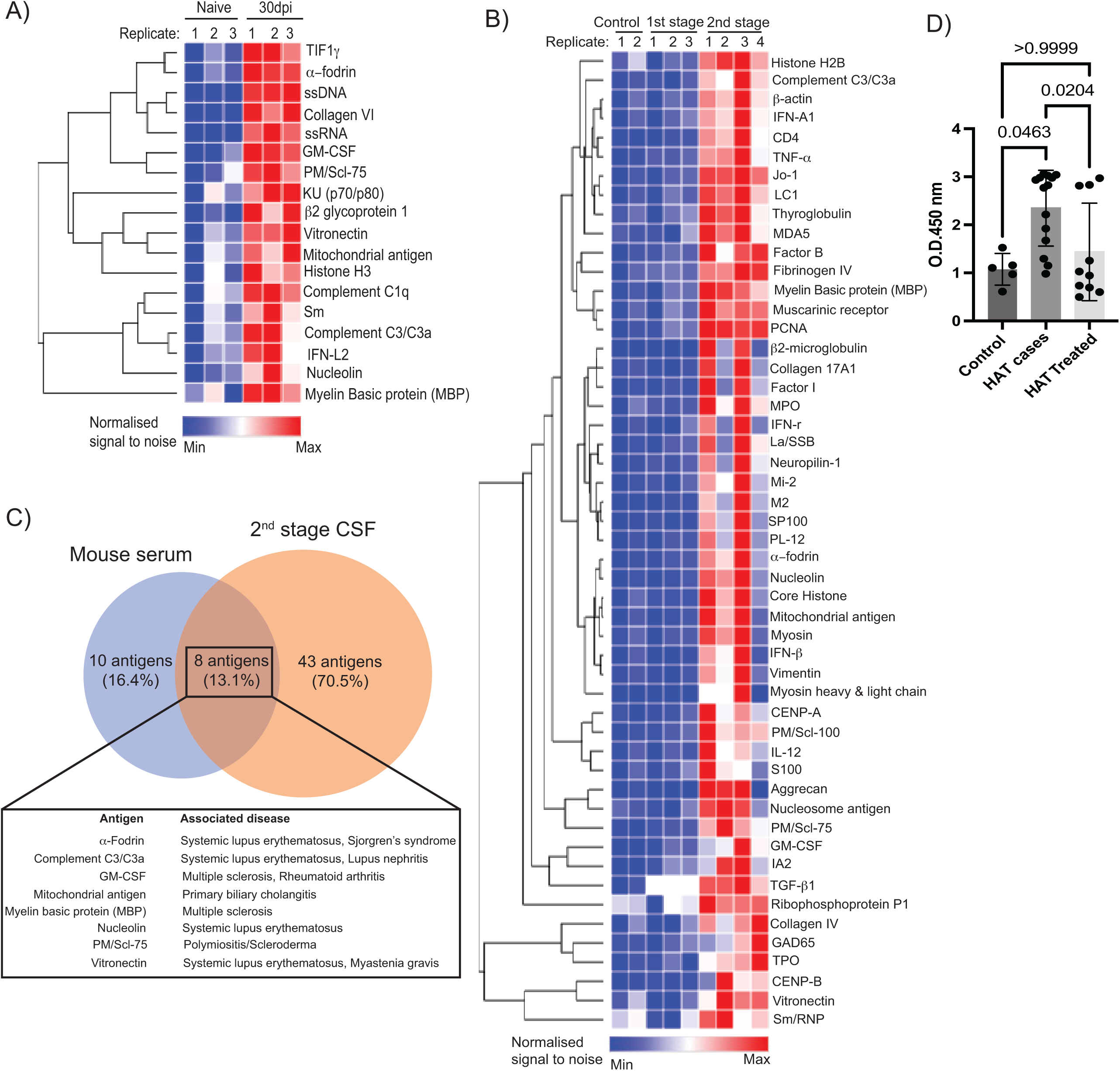
Targeted antigen screening identified shared host antigens detected by autoreactive antibodies in mouse serum and human CSF in response to *T. brucei* infection. **A)** Heatmap depicting the normalised fluorescent signal-to-noise ratio for 18 antigens significantly detected by mouse serum from infected animals at 30dpi (*n* = 3 mice) compared to naïve controls (*n* = 3 mice). The selected genes were chosen based on significant level in pairwise comparisons between naïve and infected samples using a parametric two-sided *T* test. Pairwise comparisons resulting in a *p* value < 0.05 were considered to be significant. **B)** As in (A) but depicting a total of 51 antigens significantly and exclusively detected in the cerebrospinal fluid (CSF) from 2^nd^ stage sleeping sickness patients (*n* = 4 patients) compared to both 1^st^ stage sleeping sickness patients (*n* = 3 patients) and healthy donors (*n* = 2 donors). **C)** Ven diagram depicting host antigens identified in this screening that were commonly detected by autoreactive IgG antibodies in both mouse serum and human CSF, as well as those antigens that showed specie-specific responses. A table summarising the common host antigens and the disease they are often associated with is also included. **D)** ELISA analysis to examine the presence of anti-MBP IgG autoantibodies in human serum from patients with an active *T. brucei* gambiense infection (“cases”) or post-treatment (“treated”), as well as healthy African controls (“controls”). A parametric ANOVA test with multiple comparison was used to estimate statistically significant pairwise comparisons. A *p* value < 0.05 was considered significant.

## The accumulation of meningeal GL7^+^ CD95^+^ GC-like B cells and autoreactive antibodies depends upon parasite accumulation in the CNS

Our data so far demonstrate that chronic *T. brucei* infection results in the accumulation of autoreactive B cells that display a GL7^+^ CD95^+^ GC-like phenotype, likely resulting in the generation of autoreactive antibodies and subsequent local IgG deposition in the brain. We also identified MBP, a highly abundant CNS protein, to be one of the host antigens recognised by these infection-induced IgG autoantibodies in both mice and humans during chronic infections, potentially explaining the local antibody deposition observed in the brain in our histological analyses. Given that at least 30% of 2^nd^ stage sleeping sickness patients displayed elevated levels of anti-MBP autoantibodies in circulation post-treatment, likely as a result of treatment failure, we next decided to explore whether suramin treatment, used in experimental infections to clear *T. brucei* infections^18,80^, prevented the accumulation of GL7^+^ CD95^+^ GC-like phenotype and IgG deposition in the brain. In other words, whether an active CNS colonisation is necessary to trigger local B cell responses. We tried several treatment strategies based on recent studies^18,80^, the majority of which resulted in mice relapsing to the infection. This was particularly evident when treatment was started after 14dpi. In our hands, the most effective treatment regime consisted of three consecutive doses of suramin (20 mg/kg) i.p. at 5, 6, and 7dpi, consistent with previous studies^80^ (**Figure 10A**). Using this model, we observed ∼50% of mice relapsing and ∼50% of the animals completely clearing the disease, as determined by qPCR against the *T. brucei*-specific gene *Pfr2* used here as a proxy to quantify parasite tissue burden, alongside immunohistochemistry staining against the *T. brucei*-specific antigen BiP (**Figure 10B and 10C**). Interestingly, in the relapsing animals, we noted a significantly higher parasite burden in the brain compared to infected but untreated controls. Using flow cytometry, we detected a significant expansion of GL7^+^ CD95^+^ GC-like B cells in the meninges of infected animals that remained high in the relapsing animals (**Figure 10D and 10E**). However, in cured mice, the frequency of GL7^+^ CD95^+^ GC-like B cells in the meninges returned to basal levels (**Figure 10D and 10E**). Furthermore, we observed a reduction in the IgG deposition in the brain of cured mice (**Figure 10F**), reduced serum antibody titres of anti-brain IgG autoantibodies in cured mice compared to infected or relapsing animals (**Figure 10G**), and less cortical demyelination in cured mice compared to infected and relapsing animals (**Figure 10H and 10I**), indicating that an active CNS infection is required to induce the pathological antibody responses and cortical demyelination observed in response to chronic infection. However, it is worth noting that cured mice still showed signs of antibody deposition and serum levels of anti-brain IgG autoantibodies, albeit to a lesser extent to infected or relapsing animals. Taken together, our results suggest that the presence of parasites in the CNS either directly or indirectly promotes the expansion of meningeal GL7^+^ CD95^+^ GC-like B cells and antibody deposition in the brain during chronic infection.

**Figure 10.**
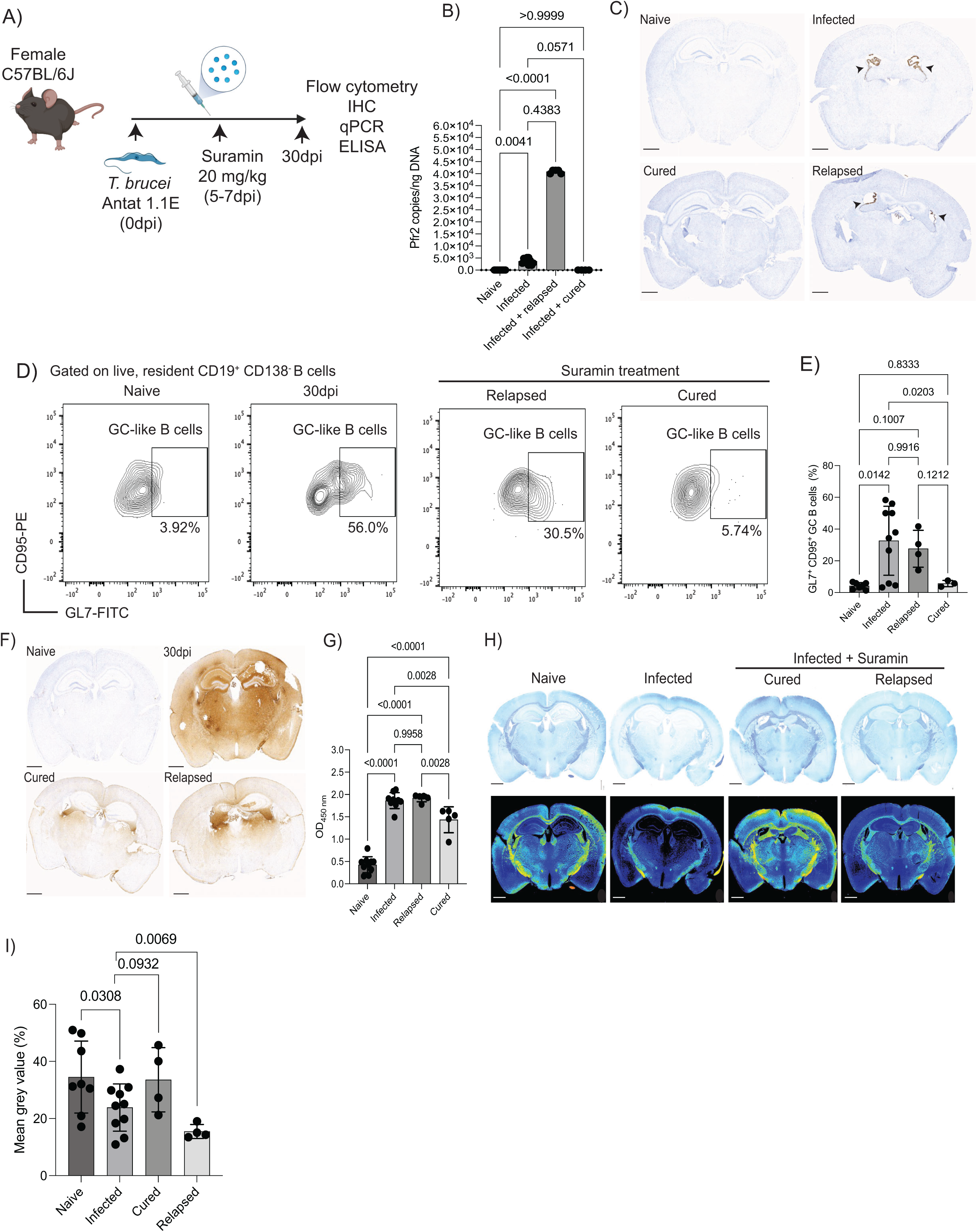
**Suramin treatment prevents the expansion of GL7^+^ CD95^+^ GC-like B cells and the IgG deposition in the mouse brain**. **A)** Overview of the experimental approach applied to prevent the CNS stage of the disease using suramin. **B)** Estimation of *T. brucei* burden in the murine brain using RT-PCR analysis to detect the parasite-specific *Pfr2* gene in naïve brain specimens and infected but untreated animals (*n* = 5 mice/group), as well as cured (*n* = 4 mice) and relapsing animals (*n* = 4 mice). These data are representative of two independent experiments. A parametric ANOVA test with multiple comparison was used to estimate statistically significant pairwise comparisons. A *p* value < 0.05 was considered significant. **C)** Immunohistochemistry staining of the *T. brucei*-specific protein BiP in brain sections from the same experimental groups as in (B). **D)** Representative flow cytometry analysis of GL7^+^ CD95^+^ GC-like CD19^+^ B cells in the murine meninges from the same experimental groups as in (B). The gating strategy to identify meningeal B cells is shown in S2C figure. **E)** Quantification of GL7^+^ CD95^+^ GC-like CD19^+^ B cells in the murine meninges from the same groups as in (B). A parametric ANOVA test with multiple comparison was used to estimate statistically significant pairwise comparisons. A *p* value < 0.05 was considered significant. **F)** Representative immunohistochemistry micrographs comparing IgG^+^ deposition in brains from naïve mice and infected but untreated mice, as well as cured and relapsed mice post suramin-treatment. Images are representative from two independent experiments. Scale bar = 1 mm. **G)** ELISA test to determine serum IgG titters of mouse brain-specific autoantibodies in the experimental groups in (B). Data are representative from two independent experiments. A parametric ANOVA test with multiple comparison was used to estimate statistically significant pairwise comparisons. A *p* value < 0.05 was considered significant. **H) Upper panel**: representative Luxol fast blue (LFB) staining as a proxy to measure myelin in brain specimens from naïve and infected mice, as well as infected mice treated with suramin, including relapsed and cured animals. Lower panel: Tissue heatmap representing the mean grey value (MGV) for the LFB staining. The calibration bar was set up so that the lowest level is 0 and the maximum MGV is 255. Scale bar = 1 mm. **I)** Percentage of demyelination, calculated here as a reduction in the mean grey intensity of the LFB staining, was calculated from two independent experiments (n = 4-10 mice/experiment). Parametric pairwise comparisons using one-sided *T* test was employed to assess significance between experimental groups. A *p* value < 0.05 was considered significant.

## Discussion

Here, we set out to characterise the local immune responses taking place within the murine meninges in response to chronic infection with *T. brucei*. Our results demonstrate that the murine meninges are dynamic structures able to support a wide range of immune responses that are triggered upon infection with *T. brucei*, resulting in the acquisition of ectopic lymphoid tissue-like signatures including the development of FDC-like cells, T_FH_ cells, and GC-like B cells undergoing class switching. We also showed that during *T. brucei* infection, the murine meninges also harbour a distinctive population of autoreactive B cells that generate IgG^+^ antibodies able to bind parasite and murine brain antigens, including MBP. Furthermore, we demonstrated that the rearrangement of the meningeal stroma, as well as the accumulation of autoreactive B cells, depend upon LTβR signalling, consistent with its lymphoid tissue properties. Lastly, we demonstrated the presence of IgG^+^ antibodies in the CSF of 2^nd^ stage gambiense HAT patients (when the parasites accumulate in the meninges and CNS) able to recognise human brain lysates and MBP, indicating that the observations using experimental infections are likely to be conserved in humans.

Focussing on the stroma, we identified several populations of meningeal fibroblasts, mostly derived from the dura mater layer of the meninges that express *bona fide* markers of mesenchymal precursor cells (*Ly6a^+^*). Interestingly, this population of meningeal fibroblasts shares many transcriptional features with omental *Aldh1a2^+^* FRCs that are known to play a critical role in modulating lymphocyte recruitment and local immune responses in the peritoneum^53^. It is tempting to speculate that there exists a pre-defined populations of fibroblasts with precursor capacity (e.g., Ly6a/Sca1^+^) in several body cavities, including the meninges, that are able to quickly sense and adapt to inflammatory responses to efficiently coordinate local immune responses^81–83^. In this context, it is plausible that the population of lymphoid stromal cells residing in the dura layer of the meninges sense the presence of *T. brucei* (e.g., *via* TLR or cytokine signalling) to promote local immunological responses, as recently proposed^84^. We propose that these populations meningeal *Ly6a^+^* fibroblast precursors could adapt to chronic inflammatory conditions, resulting in the development of stromal structures required to sustain long-lasting immunological responses *in situ*, including FRC– and FDC-like cells, as well as *de novo* angiogenesis. These observations are consistent with the idea that chronic neuroinflammation results in lymphangionesis in the CNS^85^. All of these populations participate in changes in the ECM during infection, including collagen and proteoglycan deposition and regulation of the ECM, highlighting the extensive ECM remodelling taking place in the meninges during chronic infection. The ontogeny of the meningeal FDC-like cells that we detected under chronic infection with *T. brucei*, and whether they play an active role in the neuroinflammatory process in this infection setting^86^, requires further investigation. Nevertheless, to our knowledge, this is the first report characterising the response of meningeal fibroblasts to chronic infections with a protozoan parasite, which has important implications for understanding how dynamic and adaptable the meningeal stroma is under chronic inflammatory processes.

Additionally, we predicted that meningeal MNPs are involved in chemotaxis and antigen presentation to CD4^+^ T cells. Consistent with recent studies characterising the dynamics of myeloid cells within the brain borders in response to *T. brucei* infection^9,10^, we identified several subsets of MNPs and conventional DCs that play a wide range of roles, from inflammatory responses to chemotaxis and antigenic presentation. Overall, all these myeloid subsets might offer a first line of defence against incoming parasites, and together with the meningeal stroma, might promote the recruitment and activation of adaptive immune cells required to support an efficient local immune response. It also remains to be determined whether the lymphatic structures present within the meninges play an active role in the induction of ectopic lymphoid tissues during chronic *T. brucei* infection.

We also detected an accumulation of autoreactive B cells able to produce high affinity antibodies against both *T. brucei* and the mouse brain, resembling the pathology observed under autoimmune neurological disorders including MS^87^. Previous work has demonstrated the presence of autoreactive antibodies in HAT patients and during experimental infections^88–92^, some of which are able to recognise a wide variety of heat shock proteins and ribosomal proteins^88,90^ that might be evolutionarily conserved between parasites and host. However, the local generation of autoreactive B cells in the murine meninges able to recognise murine brain lysates in response to chronic infection with *T. brucei* has not been reported before. These observations indicate that the local immunological responses in the meninges could result in the (uncontrolled) production of autoreactive antibodies, explaining their presence in the CSF of 2^nd^ stage *gambiense* HAT patients as shown in this study. Nevertheless, the mechanisms resulting in the development of local autoantibodies remains to be determined.

Based on our dataset and *in vivo* experimental approaches, we conclude that the formation of meningeal ELAs during chronic *T. brucei* infections relies on LTβR signalling. This in turn is likely to supports the expansion of meningeal FDC-like fibroblasts and the accumulation of GC-like autoreactive B cell clones and brain deposition of high-affinity antibodies, potentially aided by the local activation and development of CXCR5^+^ PD1^+^ T_FH_-like CD4^+^ T cells expressing high levels of *Il21*. In this context, our data demonstrate that chronic meningeal inflammation leads to the formation of plasma cells/FDC-like cell/CD3^+^ T cell aggregates, resembling those within the B cell follicle in lymph nodes and spleen but lacking their typical microarchitecture, to sustain the production of high affinity antibodies locally. It is important to note that the majority of the cells within the B cell compartment were assigned as IgM^+^ and IgG^+^ plasma cells as shown before^2,42,46,93,94^, although their composition (IgA^+^ vs IgM^+^ and/or IgG^+^) differs slightly from previous elegant work describing the diversity of meningeal B cells compartment during fungal infection^46^. Interestingly, in an experimental autoimmune encephalomyelitis (EAE) model, the frequency of *Igha*^+^ B cells found in the homeostatic dura mater decreased significantly followed by a significant expansion of *Ighm* and *Ighg* expression in B cells during inflammation^93^, in a process similar to the results presented in this study. These differences might be attributed to technical differences between studies (e.g., depth of coverage) or biological differences due to intrinsic differences between disease conditions. It is important to note that, in addition to the autoreactive IgG^+^ ASCs residing in the meninges and identified by ELISpot, vascular leakage (allowing the passage of IgG through the blood-brain barrier) reported in this model^95–97^ may also contribute to the IgG deposition detected within the brain. However, at present we cannot directly assess the relative contribution of each process separately. Irrespective of the routes by which autoreactive antibodies reach the meningeal barrier and/or brain parenchyma, further work is required to identify whether they arise as a result of molecular mimicry (e.g., shared epitopes between *T. brucei* and mice) or *via* bystander activation (e.g., continues TLR stimulation on B cells^98,99^). It remains unclear, however, whether the emergence of autoreactive B cells depends upon T cell-mediated responses (antigen-specific) or whether it results from T cell-independent processes (e.g., polyclonal activation, antigen-independent). In the context of T cell-dependent responses, future work is required to determine the nature of the antigens driving such specific responses, as well as the precise ontogeny of T_FH_ T cells (e.g., derived from T_H_17 T cells as recently proposed^100^) to the generation of high affinity autoreactive antibodies in the context of chronic infections remains to be delineated in more detail. It is likely that brain antigens such as MBP might be one (of several) antigen driving T cell-specific responses, in a similar process to experimental autoimmune encephalitis^101,102^.

The histological features related to antibody deposition in the meningeal and cortical spaces during chronic *T. brucei* infection are associated with cortical and white matter demyelination, which are reminiscent of the histopathological features observed in MS and other neurological autoimmune disorders^103–105^. However, it is unclear whether the cortical pathology observed in our infection model results in primary (B cells generating antibodies against myelin) or secondary demyelination, as a result of neuronal death. Importantly, *Aid*^-/-^ mice, in which B cells are unable to undergo affinity maturation, are better at controlling *T. brucei* infections due to an increase in circulating (low affinity) IgM, suggesting that class-switching might indeed be an unfavourable process for the host^106^, both in terms of parasite control and tissue immunopathology. Based on our data, local B cell affinity maturation and class-switching results in autoimmunity. In this scenario, it is tempting to speculate that most of the neuropathological features associated with chronic *T. brucei* infection derive from a disruption in peripheral tolerance resulting in maladaptive antibody responses, as recently demonstrated in variable immune deficiency patients with autoimmune manifestations^107^. Further studies are required to determine the type of antigens detected by the autoreactive antibodies generated specifically in the meninges, and to determine whether they share epitopes with *T. brucei* antigens due to molecular mimicry, as reported for EBV-induced MS^87^. Similarly, it is important to determine if B cell depletion strategies (e.g., B cell depletion approaches^108,109^, including treatment with anti-CD20 treatment^110^) or chemotherapy interventions to treat the infection ameliorate disease progression, meningeal inflammation, and cortical pathology during chronic *T. brucei* infection, similar to MS^109^. Lastly, our observations in both experimental infections and human studies indicate that sleeping sickness results in an autoimmune disorder affecting the CNS (and likely other organs), but it remains unclear whether these autoimmune disorders have an impact of sleep, contributing to the known sleep disruptions caused by this parasitic infection. In this regard, sleep disturbances are commonly reported in patients with autoimmune encephalitis^111–113^, and in narcoleptic patients^114,115^, potentially supporting a link between these pathologies. However, this remains to be further tested and the mechanisms elucidated.

Together, our data provide a novel perspective for understanding the cellular and molecular mediators that lead to the development of autoimmunity during chronic *T. brucei* infection. Furthermore, our results support the notion that the meningeal spaces are dynamic structures able to support a wide range of immunological responses, including those resulting in pathological outcomes such as autoreactive antibody deposition at the brain borders. In this context, we propose that experimental infections with African trypanosomes can be exploited to address basic questions regarding infection-induced autoimmunity and brain pathology, which could be leveraged for the treatment of complex neurological disorders of unknown aetiology such as MS in addition to the meningoencephalitic stage of sleeping sickness. Our results also highlight that chronic sleeping sickness in patients also results in the accumulation of autoreactive antibodies in the CNS, potentially driving pathology even after anti-parasitic chemotherapy. In this context, it becomes clear that a better understanding of the sequalae of the infection in human and animal health is critical but remains unsolved.

## Funding and acknowledgments

We gratefully acknowledge the contributions given by the tissue donors and their families from the Edinburgh Brain and Tissue Bank (REC 21/ES/0087). We thank Julie Galbraith and Pawel Herzyk (Glasgow Polyomics, University of Glasgow) for their technical support with library preparation and sequencing, and the technical staff at the University of Glasgow Biological Services for their assistance in maintaining optimal husbandry conditions and comfort for the animals used in this study. Similarly, we would like to thank the histopathology unit at the University of Glasgow veterinary school (In particular Frazer Bell and Lynn Stevenson) for their technical assistance with sample processing and preparation. We also thanks Professor Wendy Bailey (Liverpool School of Tropical Medicine, UK) for providing CSF samples from gambiense HAT patients from North Uganda, and Dr. Barry Bradford (Roslin Institute, UK) for his technical assistance with LFB image quantification. The authors would also like to thank the III Flow cytometry facility for their support with flow cytometry analysis, Dr. Gareth Howell (university of Manchester) for technical support with the mass cytometry experiments, and Dr. Chiara Pirillo (Beatson Institute) for technical assistance with the ELISpot assay. This work was funded by a Sir Henry Wellcome postdoctoral fellowship (221640/Z/20/Z to JFQ). AML is a Wellcome Trust Senior Research fellow (209511/Z/17/Z). PC and MCS are supported by a Wellcome Trust Senior Research fellowship (209511/Z/17/Z) awarded to AML. DB is funded by an MRC fellowship MR/V009052/1 and a Lister Institute Fellowship. NAM is supported by project Institute Strategic Programme Grant funding from the BBSRC (BBS/E/D/20002174 and BB/X010937/1). The authors declare that the research was conducted in the absence of any commercial or financial relationships that could be construed as a potential conflict of interest.

## Author contributions

**Conceptualisation:** JFQ, NAM. **Methodology:** JFQ, MCS, LKD, PC, DB, JO, GM, MAS. **Sample collection in the field:** NRKS, DMN. **Formal analysis**: JFQ, MCS, PC, MH. **Writing – original draft:** JFQ**. Writing – reviewing and editing:** JFQ, MCS, LKD, GM, DB, AM, LDL, NAM. **Funding acquisition:** JFQ. All authors approved the current version of this manuscript. The authors declare that they have no competing interests, commercial or otherwise. Correspondence and requests for materials should be addressed to Juan F. Quintana.

## Figures

**Supplementary figure 1.**
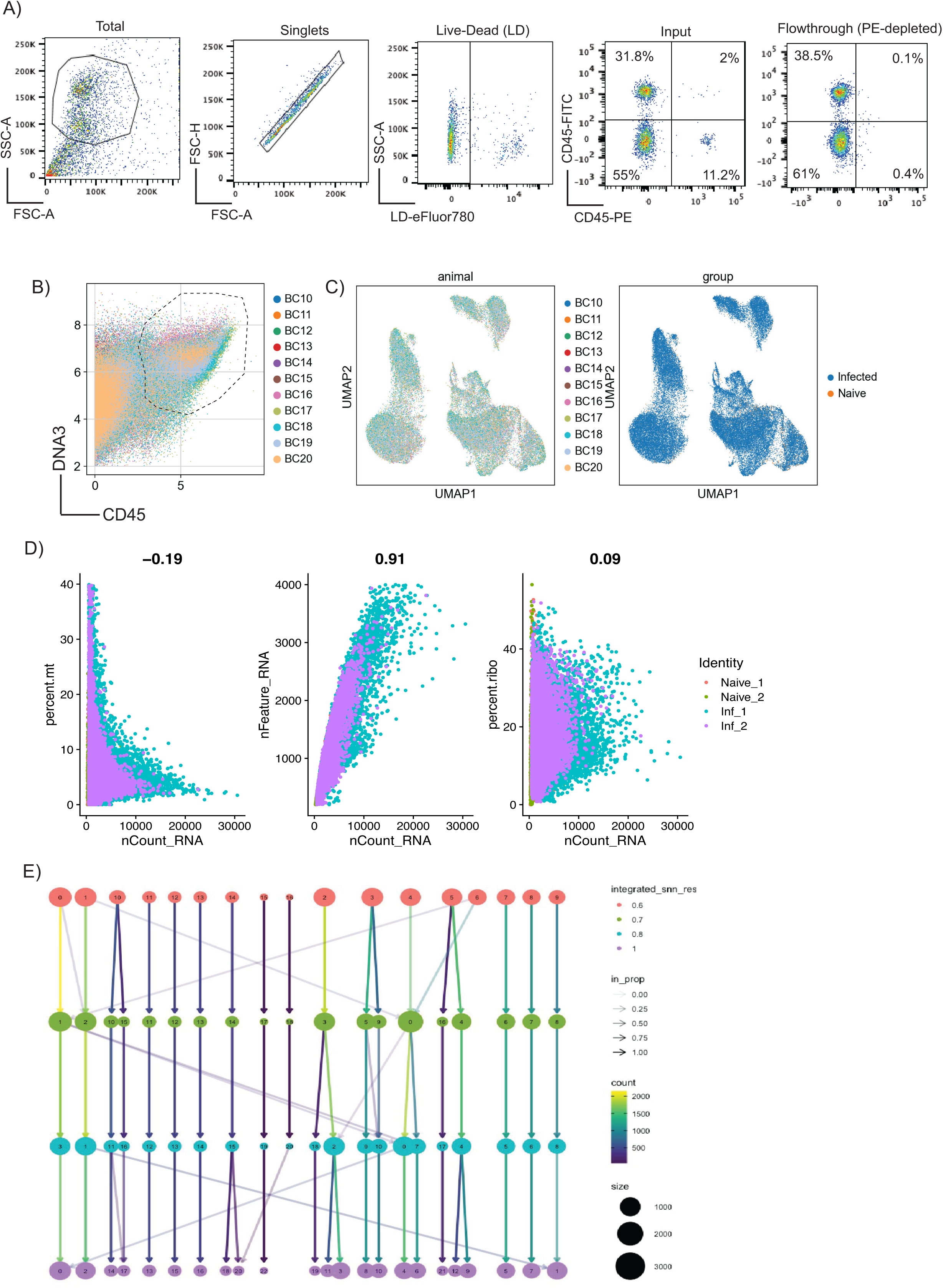
Quality control measurements of the murine single cell CyTOF and transcriptomics dataset. A) Representative flow cytometry analysis of the input and flowthrough for the removal of circulating CD45^+^ immune cells using magnetic sorting. **B)** Identification of CD45^+^ cells in the CyTOF murine meninges dataset. C) Uniform manifold approximation and projection (UMAP) of the CyTOF dataset from samples (left panel; BC10-BC15: naïve; BC11-BC20: 30dpi) experimental groups (right panel) after CD45^+^ cell clustering. **D)** Number of Unique molecular identifies (UMIs), genes, mitochondrial reads, and library complexity (Log10 UMIs/gene) after applying filtering parameters. **E)** Clustree output representing the relationship between different cell clusters at various levels of resolution using the function *FindClusters*.

**Supplementary figure 2.**
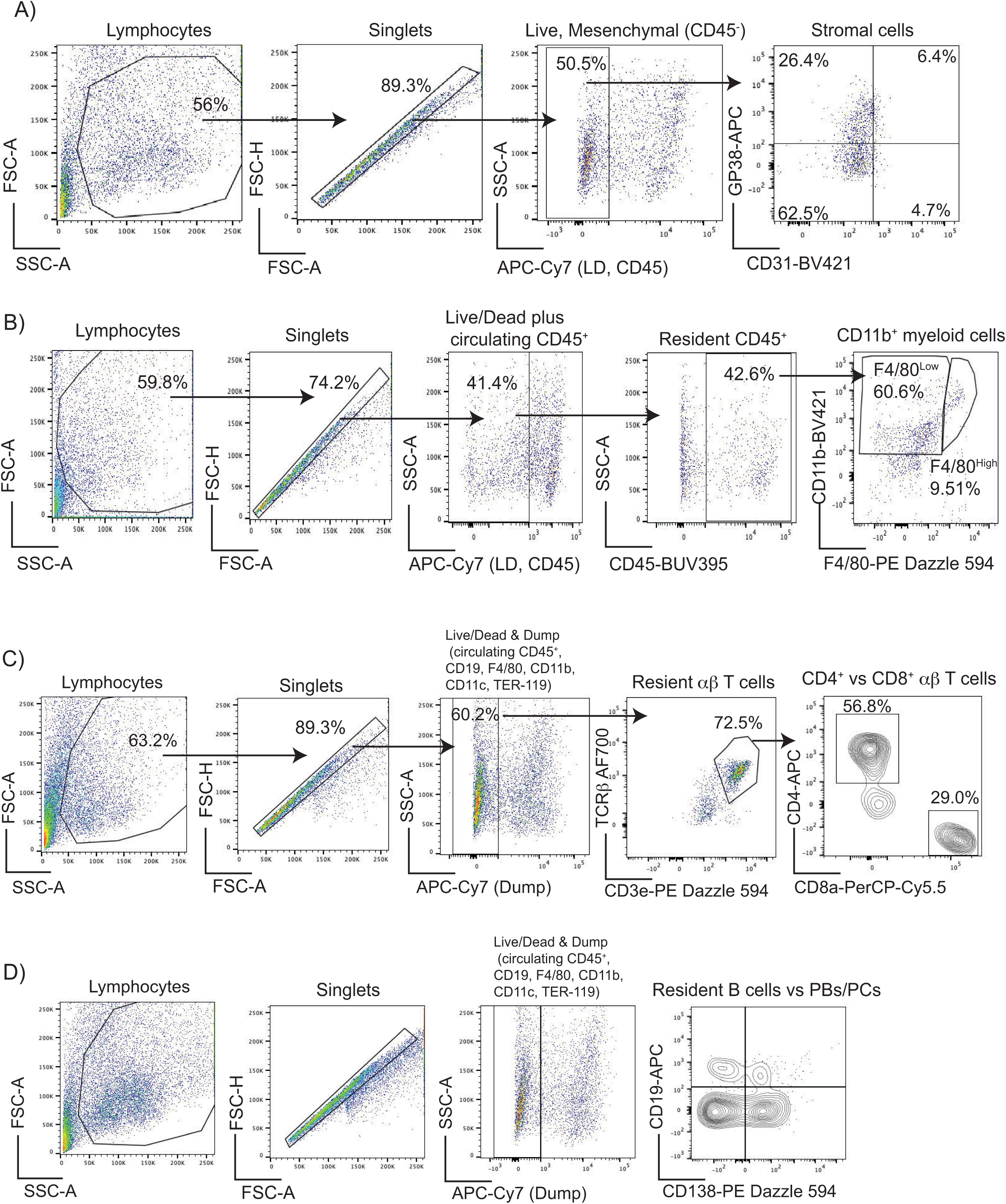
Gating strategies for flow cytometry analysis. Gating strategies to identification stromal cells (**A**), resident myeloid cells (**B**), resident CD4^+^ and CD8^+^ αβ T cells (**C**), and B cells/plasma cells (**D**).

**Supplementary figure 3.**
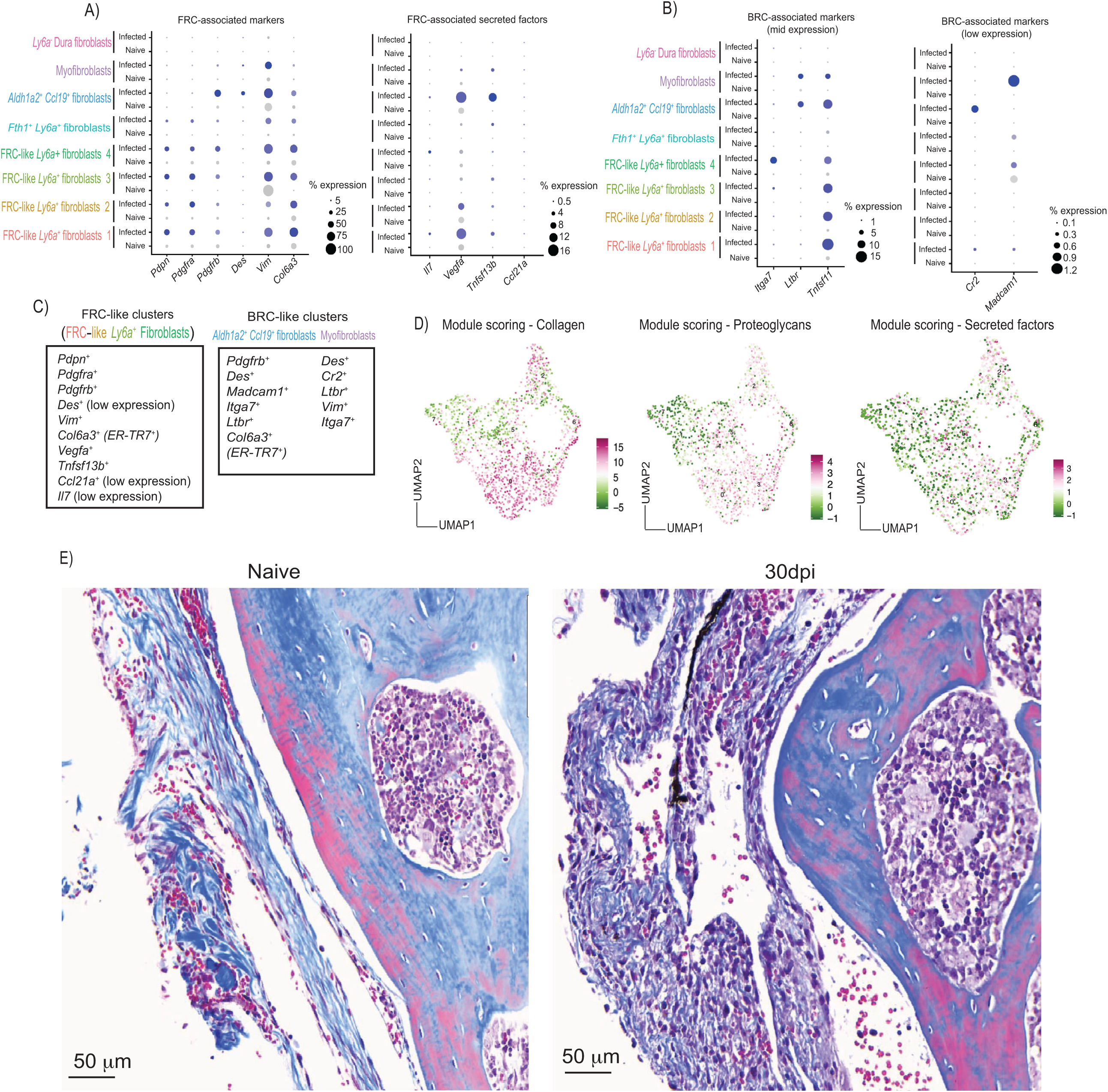
Expression of lymphoid stromal cells marker genes within the dura fibroblasts. A) Left: Expression level of Marker genes typically associated with fibroblast reticular cells (FRCs). **Right:** Expression level of genes encoding for secreted factors resealed by FRCs. **B)** Marker genes moderately (left) or lowly (right) expressed typically associated with B zone reticular cells (BRC). **C)** Broad classification of the various fibroblasts clusters as FRC– or BRC-like clusters based on the marker genes shown in (A) and (B). In all cases, the size of the dot represents the proportion of cells expressing the markers indicated in each plot. **D)** Feature plots depicting the results from module scoring of the different categories within the MatrisomeDB including collagen and proteoglycan production and deposition, and secreted factors. **E**) Masson’s trichrome staining depicting collagen deposition (blue), and keratin (pink) in sagittal skull sections containing dura meninges from naïve and infected animals. Scale bar = 50 μm.

**Supplementary figure 4.**
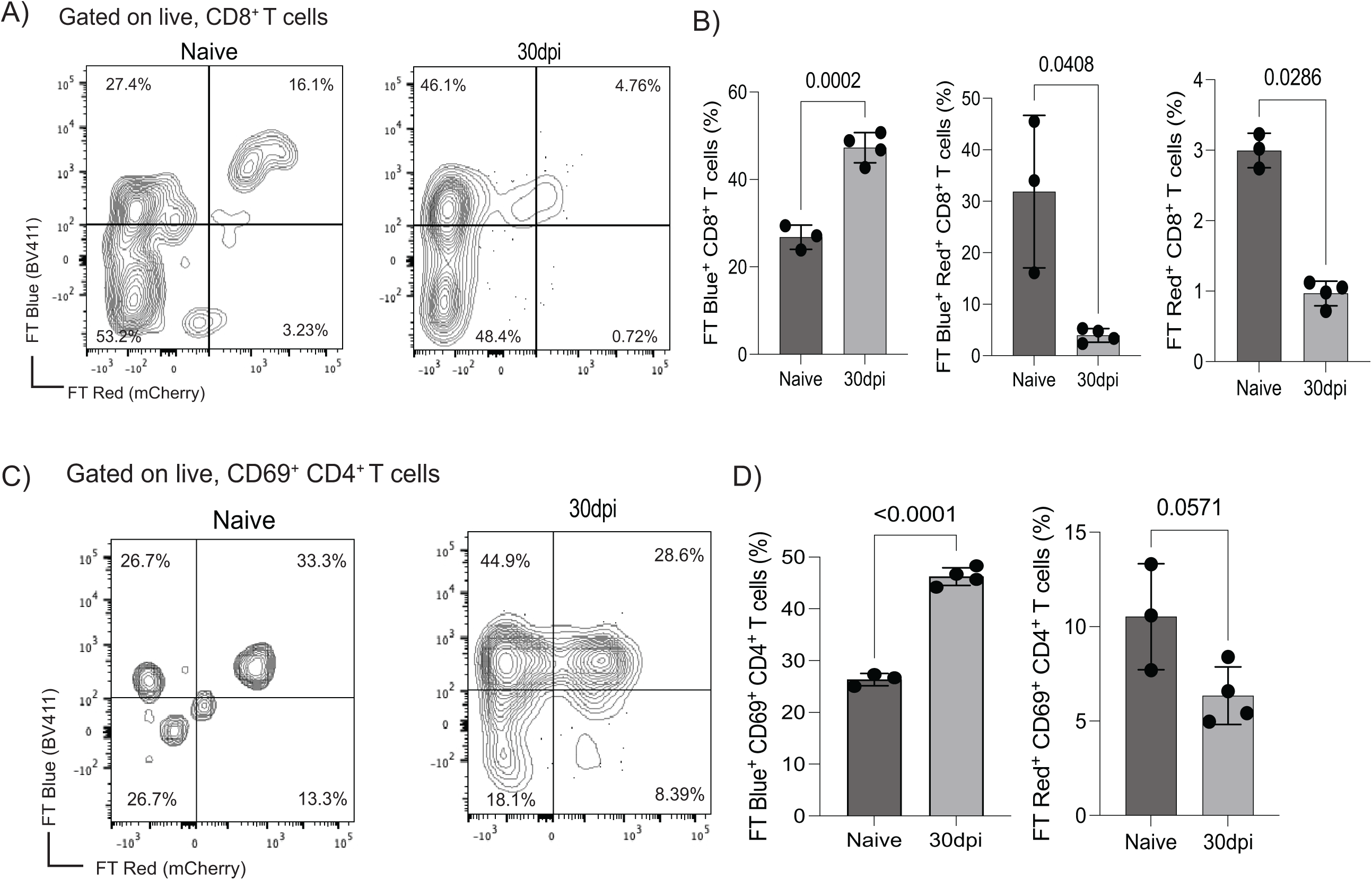
Local T cell activation in the meninges in response to T. brucei infection using the *Nur77*^Tempo^ reporter mice. A) Representative flow cytometry analysis to determine TCR engagement in CD8^+^ T cells in the *Nur77*^Tempo^ reporter mouse line. In this model, T cell activation dynamics can be discriminated between *de novo* (FT blue^+^) versus historical (FT red^+^) MHC-dependent TCR engagement. **B)** Quantification of the flow cytometry data from **(A)** focusing exclusively on FT blue^+^ or FT red^+^ CD8^+^ T cells. Data points indicate biological replicates for each panel and are representative from two independent experiments. A two-sided, parametric T test was employed to assess significance between experimental groups. A *p* value < 0.05 was considered significant. **C)** Representative flow cytometry analysis to determine TCR engagement in CD69^+^ CD4^+^ T cells in the *Nur77*^Tempo^ reporter mouse line. **D)** Quantification of the flow cytometry data from **(A)** focusing exclusively on FT blue^+^ or FT red^+^ CD69^+^ CD4^+^ T cells. Data points indicate biological replicates for each panel and are representative from two independent experiments. A two-sided, parametric T test was employed to assess significance between experimental groups. A *p* value < 0.05 was considered significant.

**Supplementary figure 5.**
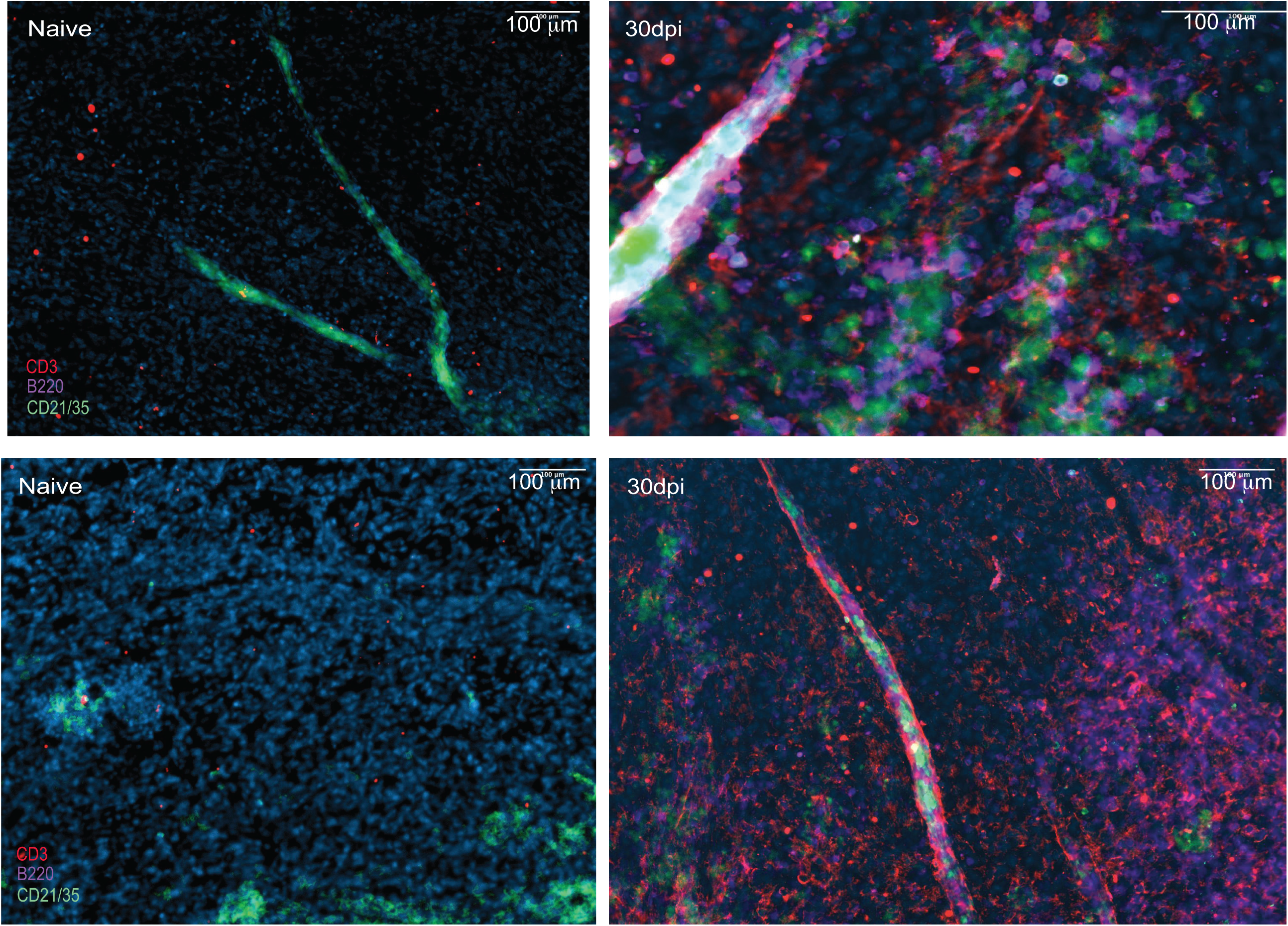
Meningeal lymphoid-like aggregates in response to *T. brucei* infection. Additional imaging analysis of whole-mount meninges from naïve (left) and infected (right) of CD21/CD35^+^ follicular dendritic cells (green), as well as CD3d^+^ T cells (red) and B220^+^ B cells (purple). The data correspond to data obtained from two independent meningeal preparations; Replicate 1 and 2 shown in top and bottom panels, respectively. DAPI was included as nuclear staining. Scale = 100 μm.

**Supplementary figure 6.**
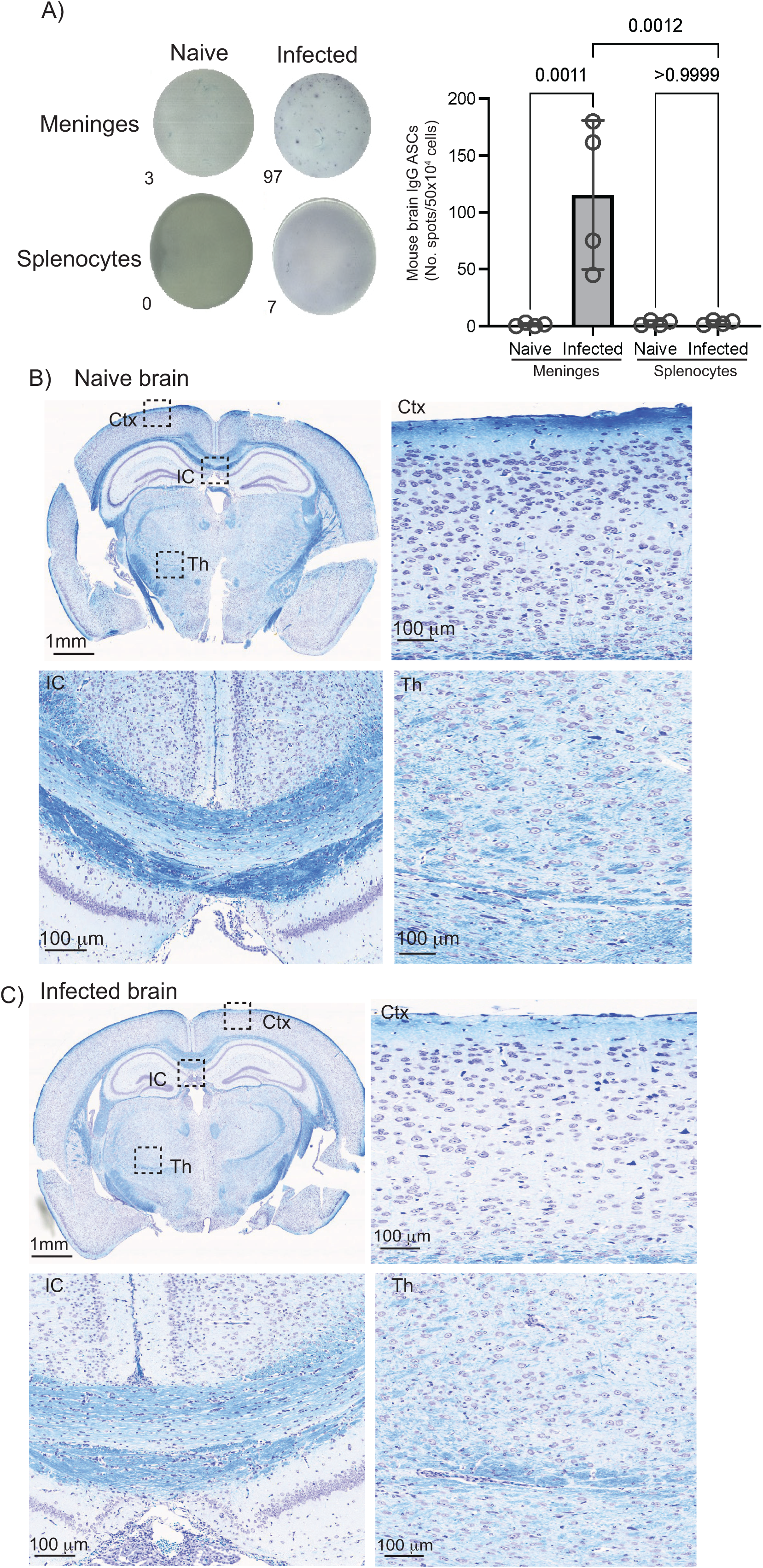
Chronic *T. brucei* infection induces demyelination in the CNS. A) Left: Representative ELISpot images depicting mouse brain-specific IgG^+^ ASCs from naïve and infected murine meninges and splenocytes after 30dpi with *T. brucei*. **Right:** Quantification of ELISpot results from mouse brain-specific IgG^+^ ASCs in meningeal preps and splenocytes (*n* = 4 mice/group). A *p* value < 0.05 is considered significant. Luxol Fast blue staining to determine myelination in naïve (**B**) and infected (**C**) coronal brain sections. Insets how areas of the cortex (Ctx), internal capsule (IC), and thalamus (Th).

**Supplementary figure 7.**
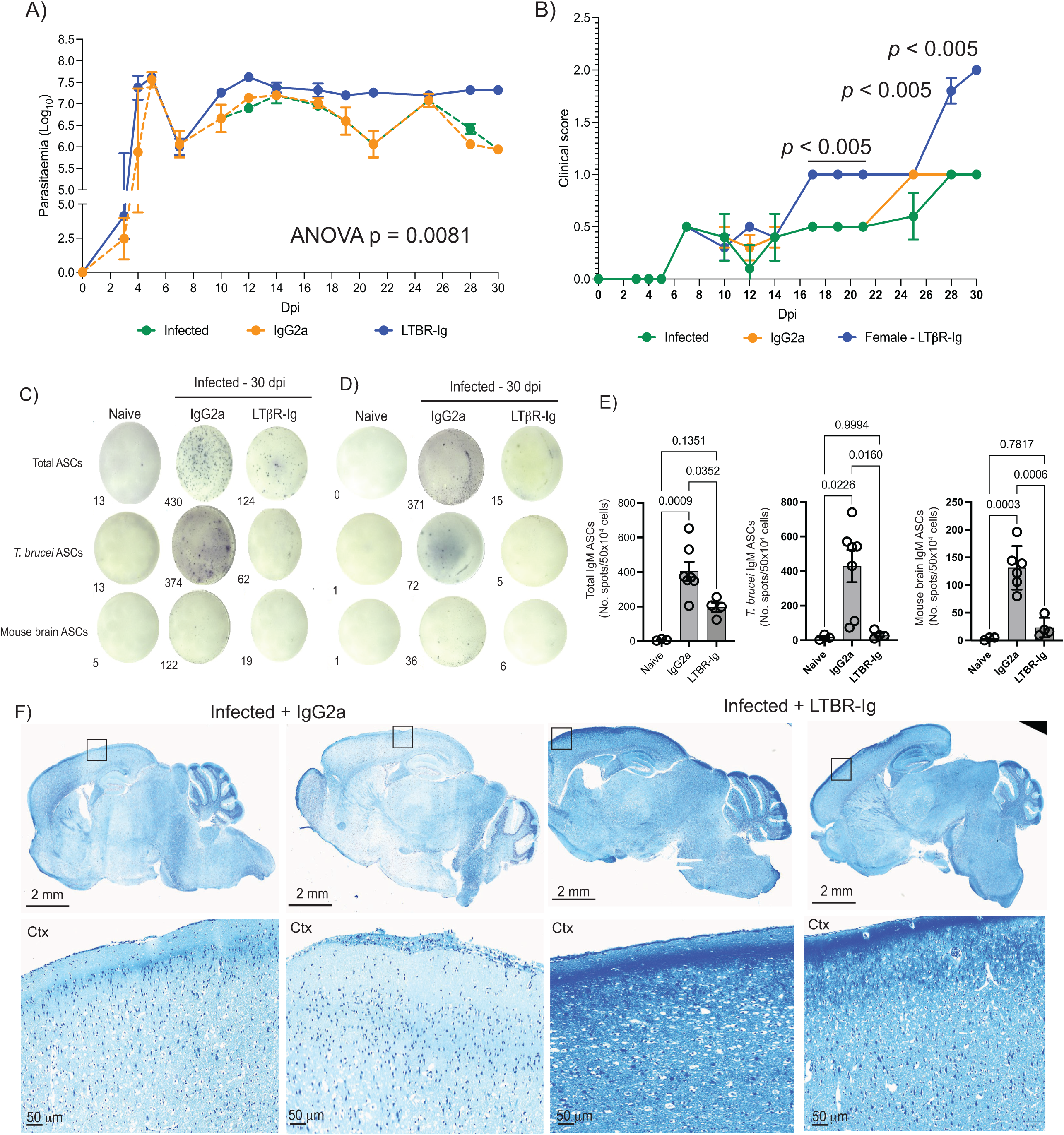
LTβR signalling blockade results in uncontrolled parasitaemia and IgM^+^ B cell accumulation in the meninges. Parasitaemia (**A**) and clinical scoring (**B**) of *T. brucei*-infected mice treated with the LTβR-Ig fusion protein (blue line). *T. brucei*-infected mice alone (green line) or infected mice treated with an irrelevant IgG2a antibody (orange line) were used as controls. For parasitaemia, an ANOVA test with multiple comparisons was conducted. For clinical scoring, pairwise comparisons were conducted using a non-parametric *T* test. In all cases, a *p* value < 0.05 was considered significant. Representative ELISpot results for meningeal IgM^+^ (**C**) and IgG^+^ (**D**) antibody secreting cells (ASCs), including total ASCs (top panel), *T. brucei*-specific ASCs (middle panel), and mouse brain-specific ASCs (bottom panel). The number of spots detected by the automated analysis software is also included. **E**) Quantification of ELISpot results including total IgM^+^ (left panel), *T. brucei*-specific IgM^+^ (middle panel), and mouse brain-specific IgM^+^ antibody secreting cells (right panel) in naïve mice, mice treated with an irrelevant IgG2a antibody, and mice treated with LTβR-Ig (*n* = 4 – 9 mice/group). A *p* value < 0.05 was considered significant. F) Luxol Fast blue (LFB) staining to determine myelination in sagittal brain sections from infected mice treated with LTBR-Ig or with an irrelevant IgG2a antibody control naïve (2 replicates per condition). Insets how selected cortical areas. Ctx, Cortex. Scale bar: 1mm (whole image) or 50 μm (insets).

**Supplementary figure 8.**
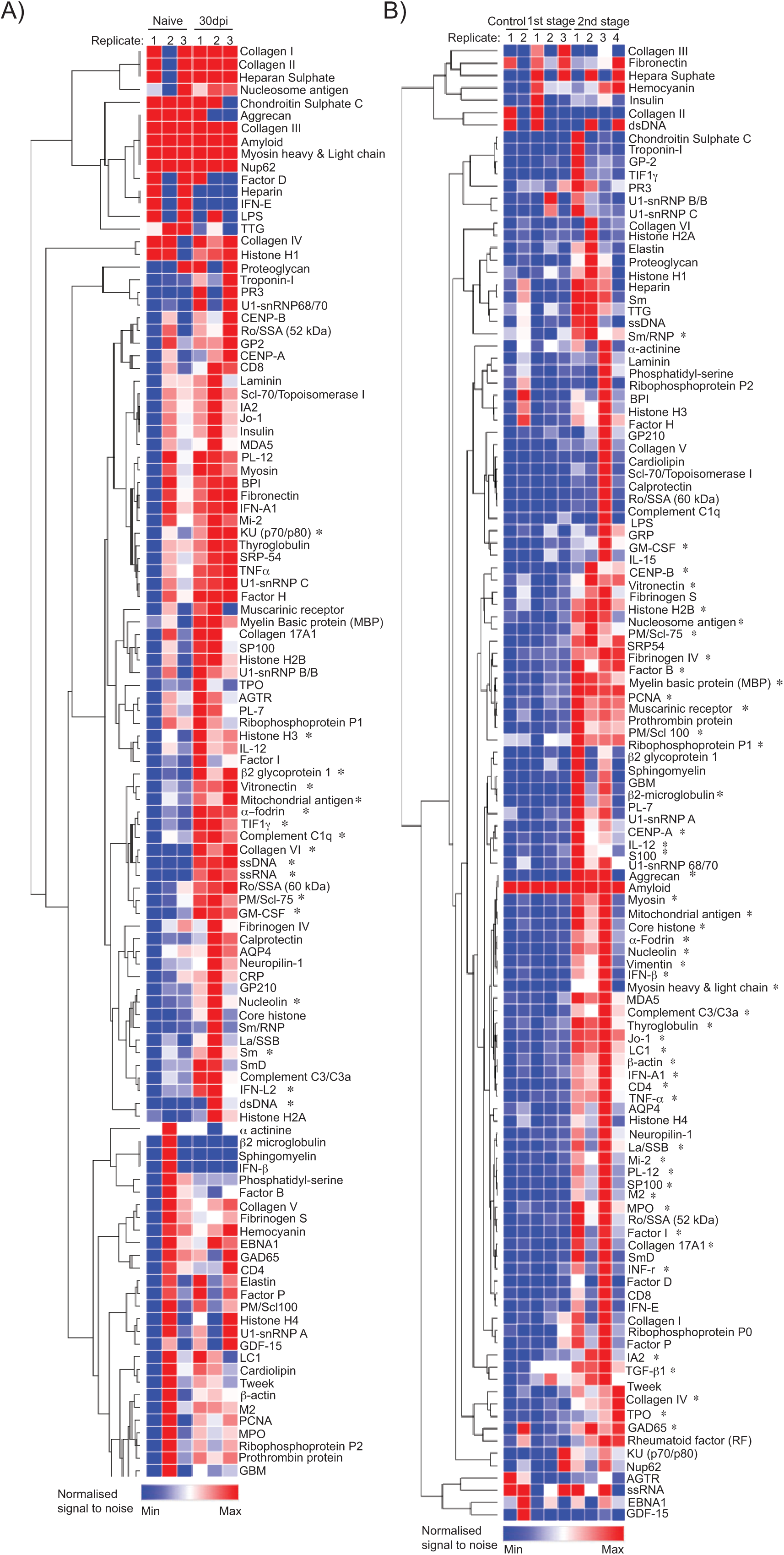
Targeted antigen screening identified shared host antigens detected by autoreactive antibodies in mouse serum and human CSF in response to *T. brucei* infection. Heatmap depicting the normalised signal-to-noise ratio for IgG binding against a panel of 120 host antigens. The data was acquired from naïve and infected murine serum (**A**) at 30dpi (*n* = 3 mice/group), and human CSF (B) from healthy donors (*n* = 2 donors), 1^st^ stage sleeping sickness patients (*n* = 3 samples), and 2^nd^ stage sleeping sickness patients (*n* = 4 samples). The asterisks denote samples that were statistically significant in a pairwise comparison analysis using parametric T test. A *p* value < 0.05 is considered significant.

**Table S1.** Quality control including mean reads per cell and median genes per cell before and after filtering out low quality cell types.

**Table S2.** Overview of the major cell types detected in the single cell dataset (resolution of 0.7). The total number of cells per cluster, percentages, and marker genes are also included.

**Table S3.** Marker genes identified for the cells within the fibroblast clusters obtained after subsetting (resolution = 0.6). The total number of cells per cluster, percentages, and marker genes are also included.

**Table S4.** Marker genes identified for the cells within the mononuclear phagocytes (MNP) cluster obtained after subsetting (resolution = 0.3). The total number of cells per cluster, percentages, and marker genes are also included.

**Table S5.** Marker genes identified for the cells within the T cell cluster obtained after subsetting (resolution = 0.7). The total number of cells per cluster, percentages, and marker genes are also included.

**Table S6.** Metadata associated with the human cerebrospinal fluid samples included for autoantibodies against human brain lysates by ELISA.

**Table S7.** Normalised signal-to-noise ratio (SNR) of targeted host antigen screening conducted using mouse serum and human CSF. The SNR values for each of the 120 antigens are included, as well as a parametric T test conducted to determine the level of significant in pairwise comparisons between infected mice and naïve controls, or between 2^nd^ stage sleeping sickness CSF samples and 1^st^ stage sleeping sickness or healthy donors.

